# Multi-modal Single-molecule Imaging with Continuously Controlled Spectral-resolution (CoCoS) Microscopy

**DOI:** 10.1101/2020.10.13.330910

**Authors:** Jonathan Jeffet, Yael Michaeli, Dmitry Torchinsky, Ifat Israel-Elgali, Noam Shomron, Timothy D. Craggs, Yuval Ebenstein

## Abstract

Color is a fundamental contrast mechanism in fluorescence microscopy, providing the basis for numerous imaging and spectroscopy techniques. The ever-growing need to acquire high-throughput, dynamic data from multicolor species is driving the development of optical schemes that optimize the achievable spectral, temporal, and spatial resolution needed in order to follow biological, chemical and physical processes. Here we introduce Continuously Controlled Spectral-resolution (CoCoS) microscopy, an imaging scheme that encodes color into spatial read-out in the image plane, with continuous control over the spectral resolution. The concept enables single-frame acquisition of multiple color channels, allowing simultaneous, single-molecule colocalization for barcoding and Förster resonance energy transfer (FRET) experiments. The simple control over the spectral dispersion allows switching between imaging modalities at a click of a button. We demonstrate the utility of CoCoS for multicolor localization microscopy of microRNA barcodes in clinical samples, single-molecule FRET measurements, and single-molecule spectroscopy. CoCoS may be integrated as a simple add-on to existing microscopes and will find use in applications that aim to record dynamic, multicolor localization events such as in multiplex FRET and tracking of multi-component, interacting complexes.

## Main

In recent years single-molecule fluorescence microscopy has become a vital instrument in the tool-box of biological, physical and chemical exploration^1–3^. Biological processes in general, and within the cell in particular, involve a plethora of interactions between various molecules and compounds. To understand these processes at the molecular level, one must record their dynamics and be able to distinguish different entities located within close proximity, usually at scales smaller than the resolution limit of standard optical microscopy (~250 nm). A distinct advantage of fluorescence microscopy is the ability to image and track dynamic processes with molecular specificity, allowing the resolution of multi-molecule functions and their building blocks. This specificity is achieved by tagging different molecular species with unique fluorescent reporters, enabling the discrimination of these molecules by the distinctive photo-physical characteristics of the markers. Moreover, with advances in resolving power introduced by fluorescence-based super-resolution techniques^4–7^, the characterization of biological processes at the molecular level has become routine.

Multiplexing is most commonly achieved by differentiating the markers’ spectral properties. Methods for spectral discrimination between different markers^8^ fall roughly into three categories: (i) Color channel separation by sequential switching of emission filters^9^, (ii) simultaneous multi-color imaging by color channel splitting of the imaged field of view (FOV)^10^, and (iii) simultaneous localization and spectral imaging by insertion of a dispersive element in one light path of a split FOV^11–14^.

While sequential channel imaging is simple to assemble and allows the largest field of view (FOV) of up to 120⨉120 µm^2^ ^9^, it comes at the cost of both linearly increasing acquisition time with each additional spectral channel, and the loss of temporal synchronization between those channels. The latter limitation prohibits any multicolor registration of dynamic processes with time scales shorter than the filter switching such as in single-molecule FRET (smFRET)^15^ measurements and multicolor tracking of many biological and physical processes.

Multicolor splitting of the FOV overcomes synchronization limitations and maintains the simultaneous acquisition of discrete color channels, but compromises throughput due to the reduced FOV that falls off reciprocally with the number of channels. To address these problems spectral super-resolution microscopy was recently introduced by multiple groups^11–14^. Generally, these systems split the FOV to a diffraction limited localization channel and a spectral channel in which dispersive elements convert the spectra into spatial intensity distributions. Nevertheless, these methods divide the total number of emission photons between the channels resulting in reduced localization and spectral accuracy^16^ and FOV sizes of 100-400 µm^2^ ^11,13,14,17,18^, 40 to 144 fold smaller compared to sequential imaging. A table summarizing the pros and cons of existing configurations is presented in supplementary Table S1.

An important aspect of all these methods is that higher spectral resolution inevitably leads to poorer spatial resolution and throughput. The signal to noise ratio (SNR) decreases with increased spectral resolution^16^ as photons are spread over more pixels, each contributing to the readout noise. Furthermore, increasing spectral resolution results in loss of camera real-estate, reducing overall throughput. While for super-resolved color discrimination the best SNR is provided by minimal dispersion, a more dispersive element is required for spectroscopy, when full emission spectra are resolved. Thus, the ideal spectral resolution is experiment dependent. In most current single-molecule spectral systems, the dispersion is permanently set by the installed dispersive element, and variation of the spectral resolution requires re-assembly of the optical setup.

Here we present Continuously Controlled Spectral-resolution (CoCoS) microscopy for single-molecule spectral imaging, which allows continuous tuning of chromatic dispersion, thus giving full control over the spectral resolution of the system. Taking inspiration from an apparatus designed for atmospheric dispersion correctors (ADC) in earth bound telescopes^19^, we positioned two identical direct-vision prisms (also called Amici prisms20) in the emission path of an epi-fluorescence microscope. These are used to introduce continuously controlled chromatic dispersion to the image for multicolor single-molecule detection and spectral analysis. A direct-vision prism, consists of three triangular prisms cemented together, with the two outer prisms identical to one another. The compound prism is built such that light of a given wavelength passes straight through unperturbed (the incident and emergent rays remain parallel with no change to the optical axis), while light of longer and shorter wavelengths is dispersed in opposite directions relative to this axis. When a pair of identical Amici prisms are aligned on the optical axis, their dispersion will accumulate to give double the dispersion of a single prism. When these prisms are rotated about the optical axis in opposite directions by ±90°, their dispersion will completely cancel out, allowing conventional super-resolution localization with minimal photon losses. Mounting the prisms on motorized rotators which can rotate the prisms independently about the optical axis enable to continuously fine-tune the dispersion with a scaling law^21^ of:

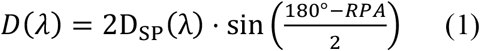

where D is the total dispersion at wavelength λ, DSP is the single prism dispersion and the relative prism angle (RPA) takes values between 0° ≤ RPA ≤ 180°, see Figure 1a for illustration.

CoCoS offers the both the flexibility to operate in different imaging modalities on a single microscope, and the ability to switch between these modalities rapidly on the same sample. Further, the CoCoS capability is easily added to existing single-molecules setups, in a simple modular upgrade. Here, we establish the principle of CoCoS, and then demonstrate three imaging modalities which correspond with key single-molecule assays: color detection, spectral analysis and smFRET.

**Figure 1.**
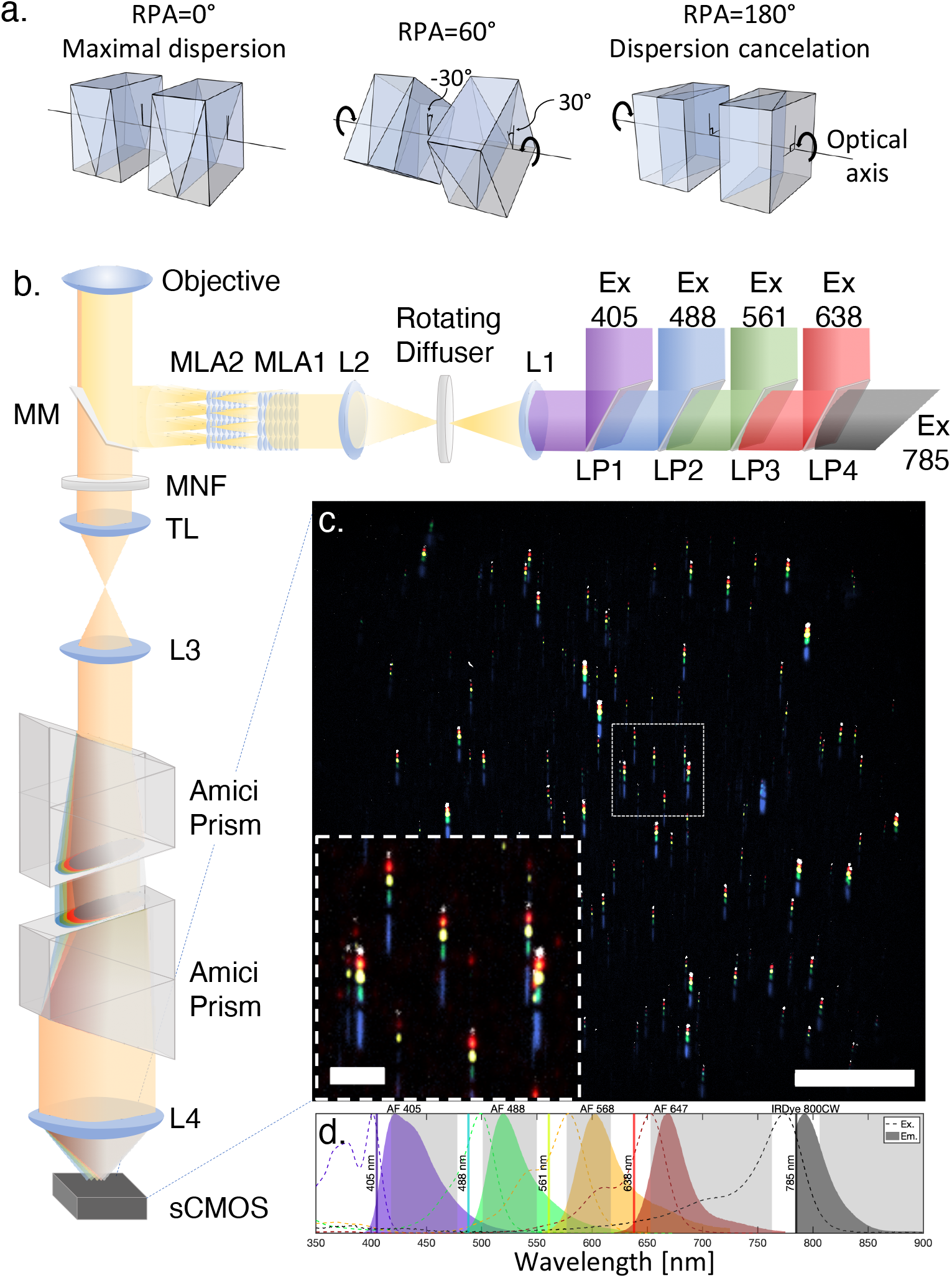
CoCoS microscopy **a**) Illustration of the prisms’ rotation about the optical axis to control the total dispersion and its axis. The relative prisms angle (RPA) about the optical axis determines the total dispersion according to equation (1), while the global angle of both prisms at RPA=0° sets the dispersion axis. **b**) Optical scheme. Five excitation lasers (Ex. 405, 488, 561, 638, 785) are combined to a single beam and reshaped to give a uniform epi-illumination profile of 130×130 µm² in the sample plane. A multichroic mirror with five spectral emission windows (MM followed by a multi-notch filter (MNF) set the emission channels of the system. The emission image is collimated out of the microscope’s tube lens (TL) by a third lens (L3) and passed through two Amici prisms located on motorized rotators. These rotating prisms allow the control of the total dispersion of the CoCoS system by setting the relative prisms angle (RPA) about the optical axis. The output beam is then focused onto a scientific CMOS camera by an imaging lens L4. **c**) Representative false color FOV of 100nm silica beads labeled with five different fluorescent dyes (Alexa Fluor (AF) 405, 488, 568, 647 and IRDye 800CW) and imaged with RPA set to 174°. Scale bar 30µm, inset 5µm. **d**) Excitation (dotted lines) and emission (solid color) spectra of the 5-dye beads, plotted against the total emission spectral channels (grey rectangles, product of MM and MNF transmission values) of the CoCoS system. Abbreviations: long-pass dichroic (LP1-4), lens (L1-4), micro-lens array (MLA1-2), multichroic mirror (MM), multi-notch filter (MNF), tube lens (TL), scientific complementary metal–oxide–semiconductor (sCMOS).

## Results

### Principle of CoCoS operation

The operational principle of the system is illustrated in Figure 1b with all technical details provided in the Methods section. Based on an inverted epifluorescence excitation scheme, we incorporated five lasers that fit the spectral windows of a five-band multichroic mirror (MM) and the matching multi-notch filter (MNF). In order to increase acquisition throughput, the combined Gaussian laser beam was reshaped to flat-top illumination^9^, providing a uniform excitation field of up to 130×130 µm^2^. The emission signal was passed through two commercial low-cost Amici prisms positioned on motorized rotators. The dispersion of the system is controlled by the rotators angle about the optical axis, permitting full computerized control over both the angle of the dispersion axis (global rotation of the two prisms), and the spectral resolution of the emission signal (controlled by the RPA). The full spectral resolution range presented in this work was scanned within 0.625 seconds with the rotators installed in our CoCoS system. However, this depends solely on the rotation speed of the rotators and can be reduced to under 17 ms with commercially available rotators. The double Amici prisms design, unlike other dispersive designs, maintains the optical axis of the system, eliminating the need for additional mirrors and significantly reduces the alignment complexity of the optical setup. Thus, the CoCoS module consisting of two rotatable prisms and a telescope (L3, L4 in Figure 1b), can be used as an add-on to any existing fluorescence imaging microscopy scheme, with no need for additional alignment.

### Continuous control of spectral resolution

We prepared a multicolor point source sample to validate the continuous dispersion control and calibrate the spectral resolution of the system. Azide functionalized, 100 nm silica beads were stained with five DBCO conjugated fluorescent dyes (Alexa Fluor 405, Alexa Fluor 488, Alexa Fluor 568, Alexa Fluor 647 and IrDye800CW) via copper-free click chemistry. The straightforward staining of the beads (any DBCO or BCN conjugated dye could be connected to the beads very efficiently) allowed for flexibility in dye selection and customization of the calibration beads to our system. Figure 1c shows a representative FOV of five-color beads with the RPA set to 174°. With this RPA, all five color windows of the beads (spectra presented in Figure 1d) are just resolvable, allowing differentiation between the colors while maintaining the highest SNR. As can be seen from the elongation of the blue channel in the figure, the fundamental dispersion of the prisms is non-linear with respect to the wavelength, with higher dispersion at lower wavelengths reaching a plateau in the near IR (see supplemental Figures S1-S3).

The continuous control over the spectral resolution allows effortless toggling between three acquisition modalities during an experiment: (i) Localization mode with RPA=180°; (ii) color detection mode, in which the different spectral channels are dispersed by the minimal resolvable amount; and (iii) full spectral mode in which the detailed emission spectra of the fluorescent dyes can be determined with sub-nm wavelength resolution. The rapid alternation between modalities allows switching with a click-of-a-button between experimental modes that previously would have required different optical setups altogether. This has many advantages, for example, in multicolor tracking^22^ or time resolved single molecule spectroscopy^23,24^, where both localization precision and spectral information are important. With CoCoS, the localization or color detection could be made first with a single frame, immediately followed by continuous spectral or spatial recording, enabling the detection of environment dependent spectral shifts or multiplex spatial dynamics (further details below).

We demonstrated the continuous control over the spectral resolution by imaging the same single silica bead labeled with four dyes (Alexa Fluor 488, Alexa Fluor 568, Alexa Fluor 647 and IrDye800CW) at various RPAs (Figure 2). The optimal spectral resolution for color detection in these beads was achieved with an RPA=174°, with the dyes just resolvable (Figure 2 – dotted red rectangle). Lower spectral resolution (higher RPA) does not allow differentiation between the dyes, while higher spectral resolution (with lower RPAs) results in the loss of throughput due to increased spreading of the molecule’s image, and decrease in SNR as the signal is spread over more pixels. Using equation (1) we calculated the optimal RPA for different commonly used dye pairs (Table 1), allowing 4 pixels between different color channels to avoid point spread functions (PSF) overlap. As is expected, dyes with more similar emission spectra require greater dispersion (lower RPA) to resolve; compare, for example, Cy3 – Atto550 (RPA=152.2°) with Cy3 – AF 647 (RPA=176.2°). When performing color detection on a multi-dye system, the lowest RPA between any dye pair in the experiment should be used.

**Table 1.** Minimal RPA values for spectral distinction of selected dye pairs. The presented RPA values were calculated for 3 (4) pixels separation between emission maxima values of the dye pairs for dyes in the same (different) spectral channel. These separation values allow differentiation between the dyes, while taking into account convolution with the emitters’ PSF (~3 pixels in diameter for our system). See methods and supporting information for additional information regarding the calculation procedure.

|  | AF<br>405 | AF<br>488 | CY<br>3 | ATTO<br>550 | AF<br>555 | AF<br>568 | AF<br>594 | CF<br>640R | AF<br>647 | CY<br>5.5 | IRDYE<br>800CW |
| --- | --- | --- | --- | --- | --- | --- | --- | --- | --- | --- | --- |
| AF 405 |  | 178.3 | 178.7 | 178.7 | 178.7 | 178.8 | 178.9 | 179.0 | 179.0 | 179.1 | 179.2 |
| AF 488 | 178.3 |  | 174.1 | 174.9 | 175.2 | 176.4 | 176.9 | 177.6 | 177.7 | 177.9 | 178.3 |
| CY 3 | 178.7 | 174.1 |  | 152.2 | 161.4 | 173.2 | 175.0 | 176.0 | 176.2 | 176.8 | 177.7 |
| ATTO 550 | 178.7 | 174.9 | 152.2 |  | 120.5 | 171.0 | 173.8 | 175.6 | 175.8 | 176.5 | 177.5 |
| AF 555 | 178.7 | 175.2 | 161.4 | 120.5 |  | 169.3 | 173.1 | 175.3 | 175.5 | 176.3 | 177.4 |
| AF 568 | 178.8 | 176.4 | 173.2 | 171.0 | 169.3 |  | 160.4 | 172.9 | 173.5 | 175.1 | 176.9 |
| AF 594 | 178.9 | 176.9 | 175.0 | 173.8 | 173.1 | 160.4 |  | 170.3 | 171.3 | 173.9 | 176.5 |
| CF 640 | 179.0 | 177.6 | 176.0 | 175.6 | 175.3 | 172.9 | 170.3 |  | 112.0 | 167.6 | 174.4 |
| AF 647 | 179.0 | 177.7 | 176.2 | 175.8 | 175.5 | 173.5 | 171.3 | 112.0 |  | 164.6 | 174.0 |
| CY 5.5 | 179.1 | 177.9 | 176.8 | 176.5 | 176.3 | 175.1 | 173.9 | 167.6 | 164.6 |  | 171.5 |
| IRDYE 800CW | 179.2 | 178.3 | 177.7 | 177.5 | 177.4 | 176.9 | 176.5 | 174.4 | 174.0 | 171.5 |  |

**Figure 2.**
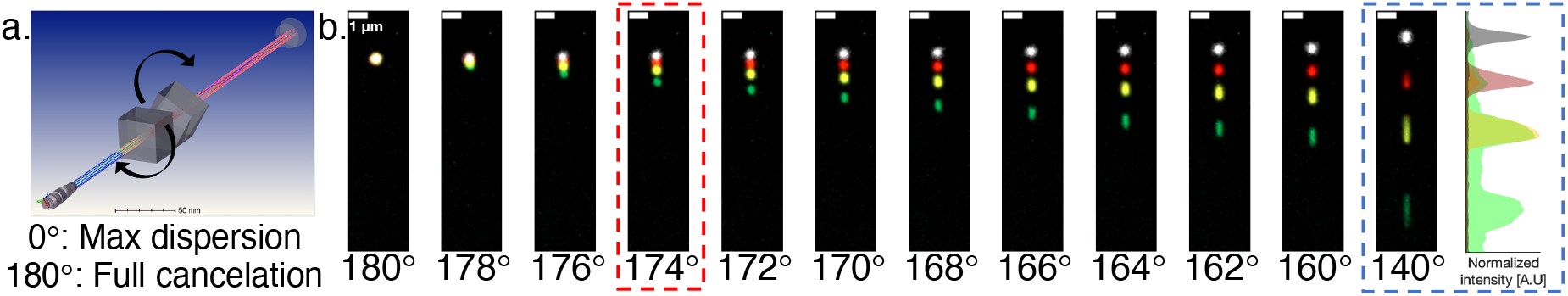
Multiple false color images of the same four-color bead consecutively excited with four excitation sources (Ex. 488, 561,638 and 785). Spectral resolution is controlled in real-time and the entire set of images was taken in under two seconds. **a**) Illustration of the prism rotation about the optical axis. **b**) The dispersion in the image plane for the stated RPAs. The motorized dispersion control allows toggling between three acquisition modalities: a-chromatic localization mode with full dispersion cancellation (180°), color detection mode with maximal SNR (for this dye combination achieved with RPA set to 174°, red dotted rectangle), and spectroscopy mode where the full spectral information within the different emission channel windows is available at the expense of lower SNR (shown at RPA of 140° for presentation purposes, blue dotted rectangle). The spectra on the far right depicts an overlay of the average intensity line plot along the Y-axis following each excitation. All images are presented with the same brightness and contrast values showing the reduction in SNR as the dispersion increases.

In the case of the four-color beads, the spectral signature of the green and orange emission is already distinct at RPA=140°, allowing us to characterize the AF488 and AF568 emission spectra (Figure 2b, blue dotted rectangle). The green spectral intensity profile (488 nm excitation) indicates high FRET from AF488 to AF568, shown by the high intensity distribution in both emission channels.

### Identifying individual microRNA barcodes in a single snapshot

To demonstrate the utility and the multiplexing advantages of our CoCoS approach we conducted microRNA barcoding experiments, achieving 4-fold reduction in acquisition time over the standard imaging protocol. MicroRNAs are short non-coding RNAs, about 20 nucleotides long, which are involved in regulation of gene expression. Sensitive, accurate and rapid diagnosis of the types and amounts of microRNAs in clinical samples are essential for advanced diagnostics and prognosis^25^. One method used to characterize multiple RNA species in a single experiment is the NanoString nCounter barcoding system^26^ in which unique fluorescent barcodes are attached to specific RNA sequences enabling expression-level detection of up to ~800 different RNA sequences. The fluorescent barcodes are composed of combinations of four colors (Alexa Fluor 488, Cy3, Alexa Fluor 594 and Alexa Fluor 647) placed at six positions separated ~500nm along a DNA capture probe. Specific color combinations report the identity of RNA molecules (the barcodes are constrained such that identical colors will never be placed in neighboring barcode positions to facilitate image analysis). Counting the various barcodes enables the determination of microRNA gene expression levels with high accuracy and sensitivity at the single molecule level. The experimental assay for barcode classification with the commercial nCounter system has been thoroughly described^26^. Briefly, the barcoded samples mixed with Tetra-speck micro-spheres (used as fiducial markers) are stretched and immobilized on specialized slides. The slides are scanned and each FOV is imaged four times with different excitations to detect the four-colored barcodes sequentially. The Tetra-speck micro-spheres are then used for image registration of the four different color channels and for chromatic aberration correction (see supplemental Figure S4). Here we performed molecular analysis on clinical microRNA samples originating from human peripheral blood mononuclear cells (PBMCs; see Materials and Methods). The NanoString nCounter system was first used to analyze the distribution of microRNAs in several samples, and the various barcodes were counted and ranked according to their abundance. We then applied CoCoS to acquire the four colors in a single frame as localized 2D spatial barcodes corresponding to each microRNA in the FOV. Since the color combinations assigned by NanoString to the various target barcodes are proprietary, we chose a subset of three samples that showed a significantly elevated level of a specific microRNA. Due to their abundance, our target microRNAs were easy to identify visually in the FOV and allowed us to assign the largest population of detected barcode to the specific microRNA (other barcode examples are presented in supplemental Figure S5). We have identified three distinct barcodes corresponding to hsa-miR-150-5p, hsa-miR-142-3p and hsa-miR-223-3p, that composed 31%, 19.3% and 18.5% of the barcodes in three different PBMCs samples respectively (the full nCounter data barcode distributions are presented in supplemental Figure S6).

Figure 3 shows three examples of these microRNA barcodes imaged on our system with RPA set to 176°.

**Figure 3.**
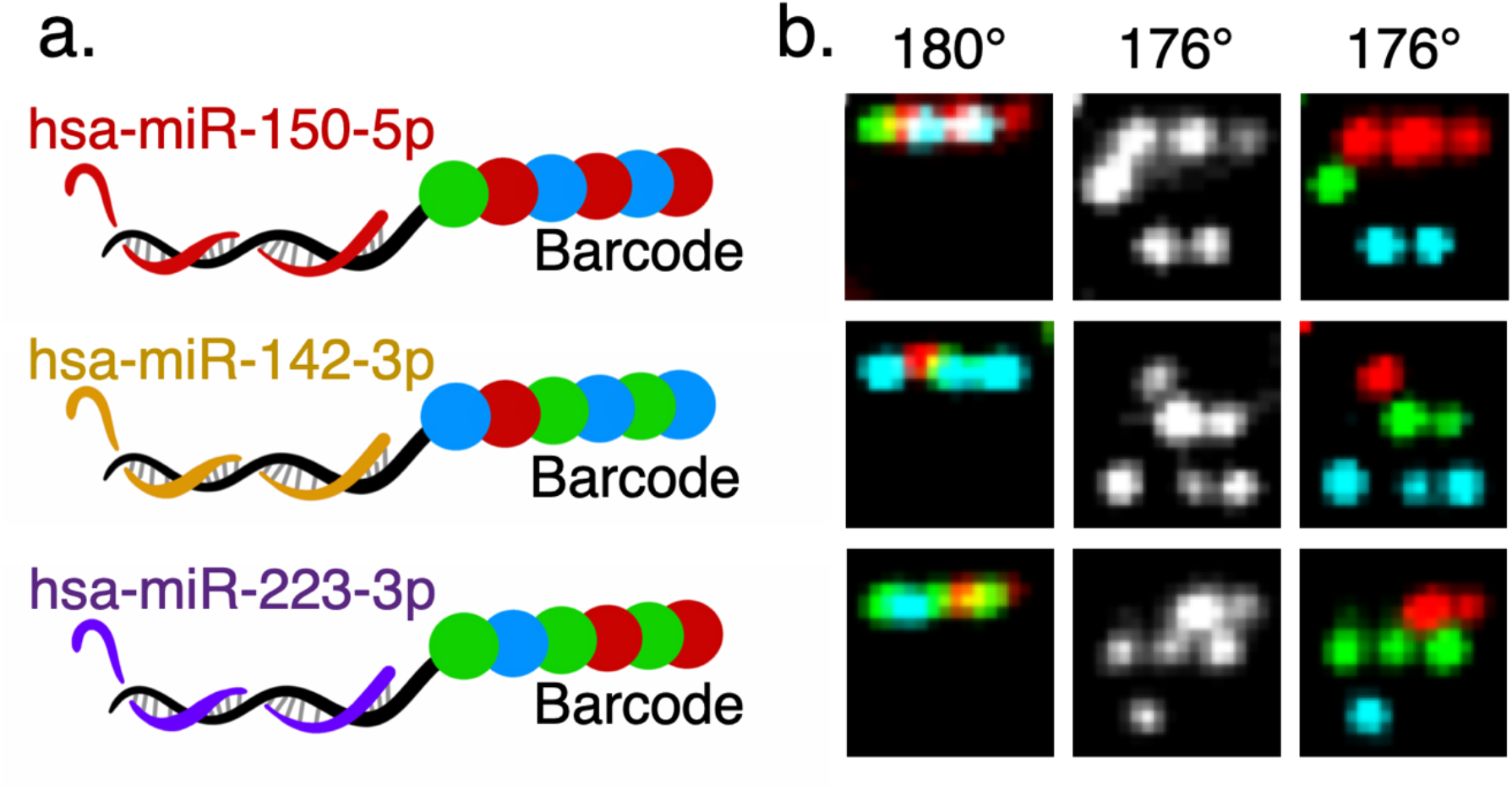
Single molecule microRNA identification using NanoString barcodes. **a**) Three types of microRNA connected to their specific reproter barcodes. **b**) False color representation of the barcodes imaged in three CoCoS acquisition modes. Left: sequential excitation with Ex. 488, 561 and 638 and with no dispersion (RPA 180°), showing the standard commercial acquisition method. Middle: a single frame acquisition with the three excitations together and RPA of 176°, displaying the ability to conver the information into a 2D single color barcode and identify the microRNA in a single frame. Right: sequencial excitation with the three sources with 176° RPA, displaying the color representation of the 2D barcode.

Operating in color detection mode, CoCoS enabled up to 4-fold increase in throughput as all the color channels are imaged simultaneously in a single snapshot, instead of four sequential acquisitions used in the nCounter system. The minimal dispersion introduced does not affect the throughput significantly, since the barcode density is usually low enough to have inter-distances larger than the dispersion (see supplemental Figure S4). By removing the time penalty for acquisition of additional colors, the combinatorial patterning of the NanoString barcodes could be expanded significantly. Every additional color to the barcode pallet contributes exponentially to the number of available combinations, for example, the addition of a fifth color will increase the number of available barcodes 5-10-fold relative to the available today. Furthermore, the simultaneous color acquisition and the fact that the dispersion creates a position shift on a single axis, eliminates the need for the channel-registration fiducial markers, allowing further increase in sample density and throughput. Furthermore, when two or more dyes per emission channel could be used, the available barcoding combinations increase 410-fold to 327,680 distinct combinations (while still avoiding the positioning of dyes from the same emission channel on adjacent positions in the barcode). This concept is demonstrated in the NanoString data, for which the spectra of two different barcoding dyes fall within the same green emission band of our system. By introducing additional dispersion with RPA of 170°, we could resolve the two dyes verifying the utility of this approach for enhanced multiplexing (see supplemental Figure S6).

### Resolving the shades of far-red fluorophores

As we show for the barcodes above, changing the dispersion to higher spectral resolutions enables the CoCoS system to distinguish between different dyes with emission spectra falling within the same emission band. Single band multiplexing enables detecting spectrally close fluorophores with a single excitation laser and reduced chromatic aberrations. Moreover, applying single band multiplexing to a multi-channel scheme significantly increases the number of distinct observables in a single experiment, dramatically expanding the available color pallet for biological experiments. We use CoCoS to extract the full spectral information and to distinguish between different markers in our far-red spectral band. Azide functionalized, 100 nm silica beads were labeled with different red dyes (CF640R, AF647, and Cyanine (Cy) 5.5) varying only slightly in their emission spectra (see supplemental Figure S7). We set an RPA of 120° according to equation (1) and Table 1 to maintain a 2-3 pixel difference between the spectral maxima of the three dyes. A spectral image (RPA=120°) was acquired together with a localization image (RPA = 180°) to determine the exact shift caused by the spectral dispersion (see Figure 4b and 4c, supplemental Figure S8 and methods section). Spectra from the different beads were then classified (as shown in Figure 4) to the various dyes using cross-correlation comparison with reference spectra obtained from single dye experiments (see supplemental Figures S9 and S10) or by comparison with theoretical spectra (supplemental Figures S11 and S12).

**Figure 4.**
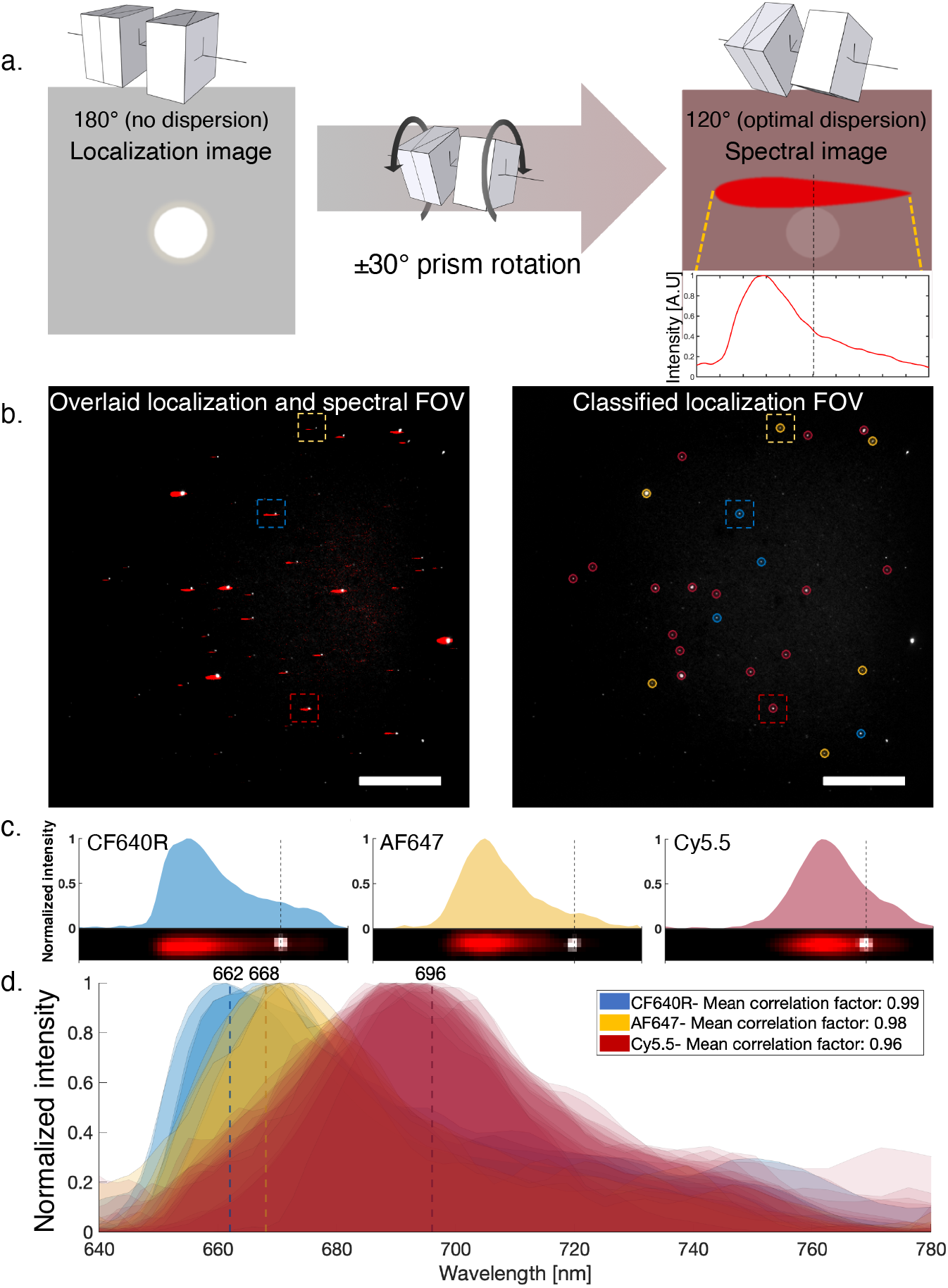
Resolving the shades of red dyes. **a)** schematic illustration explaining the procedure to obtain spectral information with CoCoS imaging. First, a localization image is captured using RPA of 180°, then the prisms are rotated to the optimal RPA for the spectral resolution of the imaged dyes (here 120°) and a second image is captured. An intensity profile of a molecule in the spectral image, with respect to the molecule’s coordinates in the localization image, contains the emission spectrum of the molecule. **b)** Three different red fluorophores were used to label 100 nm silica beads. The beads were mixed and imaged according to the procedure illustrated in a, shown in the false colored FOV image on the left. The localization image presented in white is overlaid on the spectral image in red, showing the dispersed molecular spectra. On the right, the localization image is shown after classification of the dyes according to their spectra using cross-correlation of the individual spectrum with average dye spectra (see methods). Blue, yellow and red refer to CF640R, AF647 and Cy 5.5 respectively. Colored dashed squares mark the molecules presented in panel c. Scale bars are 30 µm. **c)** Normalized spectral intensity profile of three selected molecules in panel b. Cropped images of the molecules after alignment and Gaussian filtering of the spectral image is shown at the bottom. Dashed lines indicate the localization coordinate. **d)** An overlay of all individual spectral profiles extracted from the FOV. Mean cross-correlation scores used for the classification of the spectra are shown in the legend. Vertical lines show the theoretical emission maxima of the corresponding dyes’ spectra, showing that molecular spectra with maxima differing by only ~6nm are resolvable with CoCoS.

Using this procedure, we were able to differentiate, within the same FOV, between the three dyes and classify each bead according to its spectra (see Figure 4b and 4d), showing that emission spectra with maxima differing by only 6 nm, as in the case of CF640R and AF647, are resolvable with the CoCoS system. While two consecutive frames are required to provide the reference localization to calibrate the spectra, the cost in total acquisition time is negligible considering the fast switching and potential increased multiplexing (for example, there are six, commercially available, well separated dyes suitable for our red channel alone, see supplemental Figure S13).

### High-throughput epi-fluorescence smFRET quantification

Visualizing dynamic processes with multiple colors usually results in compromised accuracy due to the time delay between sequential acquisitions in the different channels. Spectral splitting of the FOV into 2-4 spectral channels bypasses this synchronization problem but results in more complex optics and data analysis, accompanied by significant loss of camera real-estate with every additional channel. The CoCoS system enables the simultaneous recording of multiple emission channels with single molecule sensitivity, while maintaining an extremely large FOV (130×130 µm^2^). This configuration may be ideal for high-throughput single-molecule Förster energy transfer (smFRET) experiments on surface immobilized molecules. Such experiments are usually performed using total internal reflection (TIRF) illumination combined with FOV splitting, which results in a limited region for quantitative data acquisition. Here we show that our CoCoS approach allows us to perform smFRET experiments on DNA molecules under epi-illumination, removing the need for complicated TIRF optics and post-processing image registration, while gaining significant increases in throughput from our large FOV.

FRET is a non-radiative process which is sensitive to the distance between two fluorophores (a donor and acceptor^27^), and as such has been termed a ‘molecular ruler’^28^. When employed at the single-molecule level, smFRET can overcome ensemble- and time-averaging, yielding information on molecular complex formation, conformation, and dynamics (reviewed in refs.^29,30^). While these applications require the measurement of relative FRET efficiencies, methods for absolute distance measurement have now been established^15,31^ and used to determine structural models of dynamic biomolecular complexes^32–35^.

To benchmark the smFRET capability of our CoCoS system we used it to measure precise distances between two dyes in three smFRET standards, according to Hellenkamp, B. *et al*.^15^. The standards were validated in a multi-laboratory benchmark study and serve to assess the precision and accuracy of smFRET measurements. These standards are composed of short synthetic DNA molecules labeled with a donor (ATTO 550) and an acceptor (ATTO 647N) separated by 23 base pairs (bp), 15 bp and 11 bp, exhibiting low-, medium- and high-FRET efficiencies respectively (see Figure 5a). The smFRET standards were adsorbed to positively charged coverslips, and were imaged using alternating laser excitation (ALEX)^36^ with RPA=174°. At this RPA, each standard appears as a pair of point spread functions representing the donor and acceptor emissions. Multiple FOVs were recorded for each of the samples, at a frame rate of 3.3 Hz, alternating between 561 nm (donor excitation) and 638 nm (acceptor excitation). The single molecules’ emission signals were then localized, with donor emission dispersed just below the acceptor emission for each labelled DNA molecule (Figure 5b, c). Imaging the two emission signals in the same light path eliminates mismatches between channels caused by channel-specific aberrations and optical alignment. It also allows immediate real-time identification of donor-acceptor pairs without post-processing, removing the need for split-view emission channel registration. Time-traces of donor emission with donor excitation (DD), acceptor emission with donor excitation (DA) and acceptor emission with acceptor excitation (AA) were calculated with local background subtraction (Figure 5c). Using the DD, DA and AA traces, both FRET efficiency (E) and stoichiometry (S) traces were calculated per molecule and corrected using a standardized procedure^15^. (see Figure 5c for traces and supporting information for full description of the correction procedure and calculated correction factors). The corrected E and S values in each frame prior to the first bleaching event of either donor or acceptor were then plotted in a 2d histogram to show the different sample distributions (Figure 5d top panel). The ensemble averaged dye-pair FRET distances (R〈E〉) were extracted from the sample specific FRET efficiency distributions, giving R_〈E〉_= 54.3±3.8 Å, 65.8±4.6 Å, 88.4±6.2Å for the high, medium and low FRET species, respectively. These results stand in good agreement with the average experimental results from the multi-lab blind study^15^ (R〈E〉: 51.8±0.7 Å, 60.3±1.3 Å, 83.4±2.5Å for the high, medium and low FRET species, respectively). Our measurements show a slight increase in the calculated distances for all standards. This difference could be caused by the altered surface-attachment procedure applied to the DNA molecules in our experiment which were not tethered by their ends but rather adsorbed to the surface.

**Figure 5.**
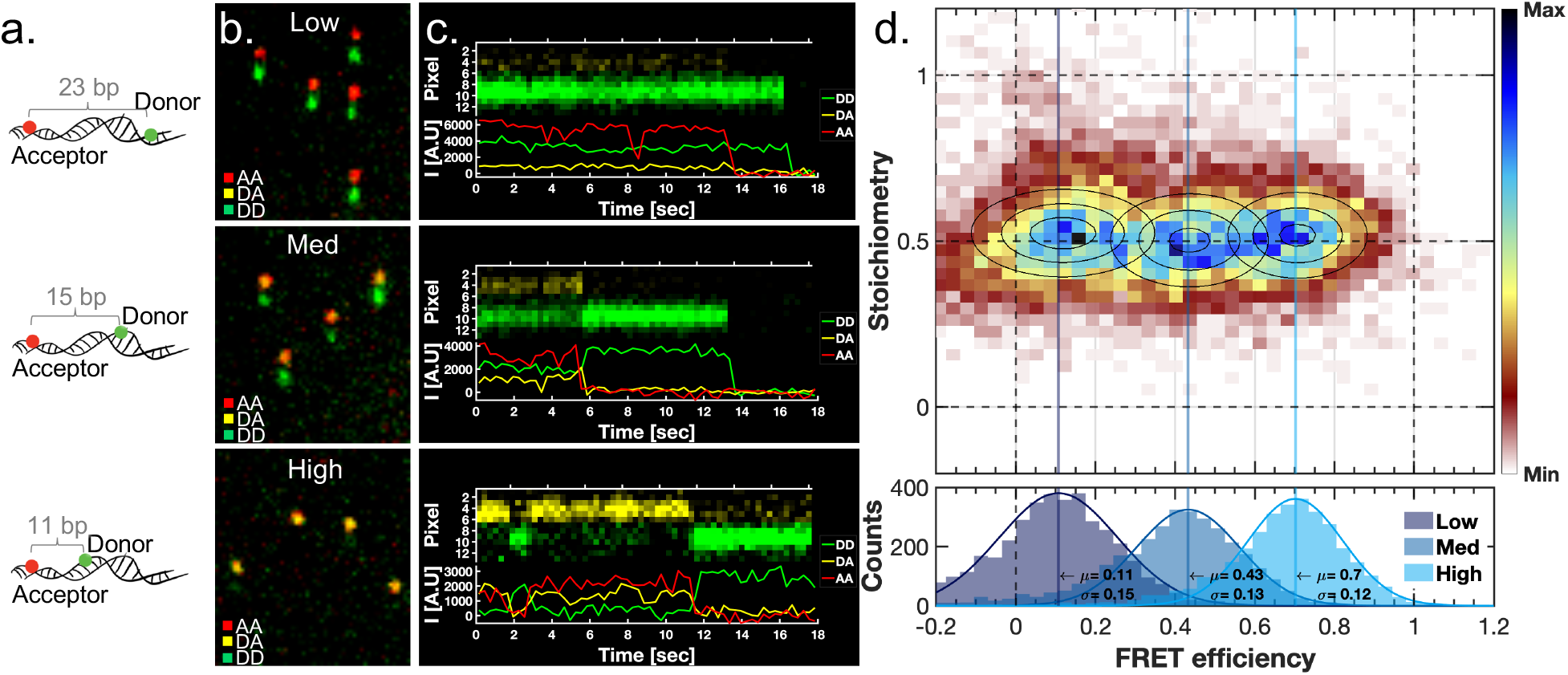
Benchmark smFRET analysis with CoCoS. **a**) Schematic drawing of the smFRET standard samples used in this experiment. Distances between donor (ATTO 550) and acceptor (ATTO 647N) are stated in base pairs (bp). **b**) Representative cropped FOVs of the different standard samples imaged with RPA of 174°. The images show false colored emission pairs in the FOV with donor excitation at 561nm (green) and acceptor excitation at 638nm (red). The acceptor emission is dispersed above the donor emission, where overlapping signals in the green and red channels represent high acceptor emission with donor excitation (i.e. high FRET appearing yellow). **c**) Representative kymographs and intensity-time traces. At the top of each panel, a false colored kymograph of a single smFRET pair emission excited with donor excitation (561nm). The acceptor’s signal (yellow) and the donor’s signal (green) are visualized in the CoCoS system one on top of the other, reducing channel registration complexity. Intensity-time traces are presented at the bottom of each panel: donor emission with donor excitation (DD-green), acceptor emission with donor excitation (DA – yellow), and acceptor emission with acceptor excitation (AA – red). **d**) E-S histogram (10722 time-points recorded from 439 molecules) of the three smFRET samples after applying correction factors and removing donor only and acceptor only time points. The fitted FRET efficiency distributions per sample (Low:142 molecules, 4098 time-points; Med:157 molecules, 3390 time-points; High: 140 molecules, 3234 time-points) are presented at the bottom panel, with their mean (µ) and standard deviation (*σ*).

The low dispersion and the lack of dichroic mirrors in the emission path, enabled us to maintain a nearly diffraction limited point spread function (PSF) for each of the colors. Thus, CoCoS provides good SNR and single molecule sensitivity even with epi-illumination, eliminating the need for the more complex and lower throughput total internal reflection fluorescence (TIRF) illumination which is the standard in the smFRET field. The CoCoS FOV is more than 6-fold larger than that of conventional dual-channel TIRF microscopes (50×50 µm²), enabling much higher throughput for smFRET measurements. In this case, only 3 FOVs per sample had to be analyzed to achieve the presented results. More importantly, since RPA of 174° is sufficient for the spectral separation of all five color channels in our system, additional FRET pairs could be easily added without any change to the setup or to the dispersion, allowing multicolor smFRET experiments with almost no reduction in throughput or time-resolution.

## Discussion

Here we introduced CoCoS microscopy, an optical add-on to wide-field epi-fluorescence microscopy, which allows continuous control over the spectral resolution of the system. A key feature of CoCoS is the ability to easily switch between different modes of operation: (i) imaging and localization, (ii) color detection / FRET and (iii) spectral analysis. With our epifluorescence microscope system we showed the ability to detect and localize up to 5 color channels simultaneously, affording up to a 5-fold reduction in multi-channel acquisition time and significantly enhancing throughput in multi-color experiments. Moreover, the number of detectable color channels is only limited by the fluorophores’ spectra and the transmission bands of the multichroic mirror and band-pass filter used in the experiment. Thus, future work can easily expand the number of possible emission channels in the color detection mode.

The key limitation of our current system stems from the prisms used in this setup. These are currently low cost, off-the-shelf prisms, which have a small cross-section that introduced optical aberrations at small RPAs due to the large field of illumination. Furthermore, the non-linear dispersion of the prisms created varying spectral resolution across the spectrum, forcing different RPA for the same spectral resolution in different wavelengths. This will be corrected in future work by carefully designing prism material and apex angles^37^.

Unlike the conventional filter-based systems, the detection of color in CoCoS is based on the spatial separation of spectral PSFs. Recent incorporation of deep-learning for PSF based color detection^38^ has shown that small variations in PSF could be exploited for color detection and differentiation between markers. With CoCoS, minute and well calibrated color-dependent PSF changes can be easily introduced, allowing full color PSF detection by deep-learning. Unlike PSF based color detection where a liquid-crystal spatial light modulator (SLM) device is used to control the PSF^38^, CoCoS has almost no photon loss making it much more suitable for low SNR settings typical for single-molecules in biological contexts. An immediate attractive application is quick decoding of the 2D spatial barcodes presented here for single-shot imaging of NanoString barcodes. The distinct PSF structures created by each color combination are ideally suited for machine-vision analysis, and may allow real-time data processing.

The ability to record emission signal from all these color channels simultaneously without splitting the FOV, opens the door for highly multiplexed smFRET experiments. Utilizing five or more donor-acceptor pairs multiplexed on the same observed complex, will allow better reconstruction of molecular structure and dynamics for proteins or other molecular assemblies^39^. This feature not only expands the current limit on multi-color smFRET which currently stands on four colors^10,29^, but also dramatically simplifies the optical setup while expanding the FOV and throughput of the system by more than 5-fold. Further expanding the color multiplexing capabilities from the discrete 5 color-channel multiplex to a continuous spectral differentiation within the same spectral band, will further boost the available fluorescent marker palette. With the current ability to quickly toggle between localization and full spectral information, both tracking and differentiating multiple single molecule markers within the same spectral window is made possible. This makes CoCoS attractive for multi-component single-molecule tracking with full spectral information. Furthermore, the available spectral recording will allow us to exploit environmental^14,23^ or voltage^24^ dependent single molecule spectral shifts.

## Methods

### Optical Setup - excitation

The excitation module was composed of five lasers (Cobolt AB, Sweden) with wavelengths spanning the visible to NIR spectrum: 405nm (MLD 405, 250mW max power), 488nm (MLD 488, 200mW max power), 561nm (Jive 561, 500mW max power), 638nm (MLD 638, 140mW max power) 785nm (NLD 785, 500mW max power). All lasers were mounted on an in-house designed heatsink which coarse aligned their beam heights. Each laser beam was passed through a clean-up filter (LL01-405-12.5, LL01-488-12.5, LL01-561-12.5, LL01-638-12.5, LL01-785-12.5, Semrock, USA) and expanded to 12.5-20x its original diameter (LB1945-A, 4×LB1157-A, 3×LB1437-A, LD2746-B, LB1901-B, Thorlabs, USA). Two motorized shutters (SH05, Thorlabs, USA) were used for modulating on/off the solid state 561nm and 785nm lasers due to long warm up times. The beams were then combined to a single beam using long pass filters LP1-LP4 shown in Figure 1b (LP1: Di03-R405-t1-25.4D, LP2: Di03-R488-t1-25.4D, LP3: Di03-R561-t1-25.4D, LP4: Di03-R635-t1-25.4D, Semrock, USA). In order to homogenize the excitation profile of the sample, the combined beam was passed through an identical setup to the one described in the work of Douglass et al.^9^. In short, the combined beam was injected into a compressing telescope (AC254-150-A-ML, AC254-050-A-ML, Thorlabs, USA) with a rotating diffuser (24-00066, Süss MicroOptics SA, Switzerland) placed ~5mm before the shared focal points of the telescope lenses (see Figure 1). A series of 6 silver mirrors (PF10-03-P01, Thorlabs, USA) was then used to align the beam into a modified microscope frame (IX81, Olympus, Japan), through two identical microlens arrays (2×MLA, 18-00201, Süss MicroOptics SA, Switzerland) separated by a distance equal to the microlenses focal length and placed inside the microscope frame. The homogenized beam was reflected onto the objective lens (UPlanXApo 60X NA1.42, Olympus, Japan) by a five-band-multichroic mirror (MM, zt405/488/561/640/785rpc, Chroma, USA). The sample was placed on top of motorized XYZ stage (MS-2000, ASI, USA) with 890nm light emitting diode (LED) based autofocus system (CRISP, ASI, USA) which enabled scanning through multiple fields of view.

### Optical Setup - Emission

The emitted light from the fluorescent samples was gathered by the same objective and transmitted through the multichroic mirror (MM) onto a standard Olympus tube lens to create an intermediate image at the exit of the microscope frame. This image was passed through a multi-notch-filter (MNF, NF03-405/488/561/635E or FF01-440/521/607/694/809-25 multi-band-filter in the smFRET experiment, Semrock, USA) combined with a 785nm notch filter (NF03-785E-25, Semrock, USA) to block all residual laser lines. Light was then directed into a magnifying telescope (Apo-Rodagon-N 50mm, and AC08-075-A-ML, Thorlabs, USA or Apo-Rodagon-N 80mm, Qioptiq GmbH, Germany in the NanoString and smFRET experiments), with two commercial direct vision prisms (117240, Equascience, France) placed within the infinity space between the lenses and mounted on two motorized rotators (8MR190-2-28, Altechna UAB, Lithuania) controlling the prisms’ angles around the optical axis. The final image was acquired on a back illuminated sCMOS camera (Prime BSI, Teledyne Photometrics, USA).

Image acquisition was coordinated using the micro-manager software^40^, controlling camera acquisition, laser excitation, XY stage location and prism rotator angles. The camera and lasers excitation were synchronized using in-house built TTL controller based on an Arduino^®^ Uno board (Arduino AG, Italy)^41^.

## Sample preparation

### Multicolor silica beads

For the labeling reaction, 5µL of 1% suspension of azide-functionalized 100 nm silica beads in aqueous solution (Si100-AZ-1, Nanocs, USA) were diluted in 20 µL doubly deionized H_2_O (DDW) and sonicated for 1 hour. Next, the beads were labeled using a copper-free click chemistry reaction with various dibenzocyclooctyl (DBCO) activated fluorescent dyes according to the experiment.

### Five color beads

A total of 0.53 mM DBCO activated dyes (AFDye 568-DBCO, Click Chemistry Tools, USA; IRDye 800CW-DBCO, LI-COR, USA; AF647-DBCO, AF488-DBCO, AF405-DBCO, Jena Bioscience, Germany) were added to the beads solution with a relative ratio of 2:1:1:1:2 corresponding to AF405:AF488:AF568:AF647:IrDye800CW. The beads and dye solution was vortexed and pipetted vigorously to homogenously distribute all dyes and left over-night at 37 °C for the click reaction. Next, ethanol-isopropanol precipitation was used to remove excess dye molecules. Isopropanol cooled to −20 °C was added to the beads and dye solution to a final volume of 1mL. The solution was centrifuged at 4 °C for 35 minutes at 18,000 rpm, creating a pellet of labeled beads at the bottom of the tube in a solution of excess dyes and isopropanol. The solution was carefully removed without disrupting the beads pellet. The same procedure was then repeated with cooled to −20 °C 100% ethanol and then with a cooled to −20°C 70%/30% ethanol/DDW solution. Finally, the labeled beads were suspended in 50 µL DDW to create a stock solution. For imaging the beads, 3µL of the stock solution were added to 20 µL TE buffer (10 mM Tris, 1 mM EDTA, pH 8) mixed with 4 µL of 2 mM DTT and placed on pre-cleaned cover glass.

### Far-red beads

For the far-red beads experiment the same procedure was followed mixing a single 75 µM DBCO/BCN activated dye (AF647-DBCO, Cy5.5-DBCO, Jena Bioscience, Germany; CF640R-BCN, Biotium, USA) with the azide-covered nano-beads to create three different stock solutions. Due to high background noise caused by unreacted dyes, which became prominent due to the high dispersion in this experiment, we added another cleaning step at the end of the previous ethanol-isopropanol precipitation with drop-dialysis on 0.05 µm pore membranes (VMWP02500, MF-Millipore™ Membrane Filter - 0.05 µm pore size, Merck, USA).

### NanoString

For the NanoString experiment a commercial slide was taken after being processed and analyzed on the nCounter system as described by the manufacturer (for example see ref ^42^). Peripheral blood mononuclear cells (PBMCs) were collected from treatment resistant depression (TRD) patients. All samples used in this study were collected with informed consent for research use, and approved by Institutional Review Boards (IRB approval 0002-14-SHA) in accordance with the declaration of Helsinki. RNA samples were extracted using TRIzol Reagent (Thermo Fisher Scientific, MA, USA) from these PBMCs and was subjected to microRNA profiling using the NanoString nCounter microRNA expression assay (NanoString Technologies, Seattle, WA, USA). Briefly, 100 ng of total RNA was used as input material. A specific DNA tag was ligated onto the 3’ end of each mature microRNA, providing an exclusive identification for each microRNA species in the sample. The resulting material was hybridized with a panel of fluorescently labeled, bar-coded reporter probes specific to the microRNA of interest. Abundances of microRNAs were quantified with the nCounter Prep Station *via* counting individual fluorescent barcodes and quantifying target microRNA molecules present in each sample. The slide was imaged on our system with no additional treatment. Each lane in the slide consisted of a different sample and presenting various microRNA expression.

### Single-molecule FRET (smFRET)

smFRET standards^15^ of high, medium and low FRET efficiencies (ATTO550/ATTO647N with 11-bp, 15-bp, 23-bp separation between donor and acceptor respectively) were each diluted to 0.5 nM in TE pH 8 buffer and placed on silane-activated glass coverslips prepared according to an established protocol^43^.

## Acquisition parameters

All samples were imaged with the same hardware as described in the optical setup section of the methods. Due to various sample background and signal to noise ratio (SNR) conditions, sample specific acquisition parameters were used and are reported herein.

### Five color beads

The image presented in Figure 1 was acquired using sequential five laser excitation with a single laser per camera frame. The laser intensities at the laser outputs were 400 mW for the 785 nm laser and 100 mW for all other lasers, 100 ms camera exposure per frame and RPA =174°.

For spectral calibration, 13 images with RPA ranging from 180° to 140° were imaged by sequential excitation of all 5 lasers. Laser intensities of 50 mW were used for all lasers and 300 ms camera exposure was used per image (see supplemental Figure S1).

### Far-red beads

All far-red bead samples (Figure 4 and supplemental Figure S8) were imaged with the same acquisition parameters: the 638nm laser at 30mW at its output, 300ms camera frame exposure and two frames per FOV, no dispersion at 180° RPA and optimal dispersion at 120° RPA. The perpendicular shift between the two frames (along the y-axis as presented in Figure 4) is due to optical aberrations of the lenses L3 and L4, together with slight misalignment of the two prisms relative to the lenses’ optical axis. At maximal dispersion values the limited clear aperture of the prisms caused additional aberrations due to higher beam divergence causing the light to deflect from the prisms’ edges.

### NanoStrings

The NanoString samples were imaged with three lasers 488 nm, 561 nm and 638 nm operating at powers of 50 mW, 30 mW and 50 mW respectively. In each FOV and for each of the two prisms positions (RPA=180° and RPA=176°), four 300 ms exposure images were taken: using all three lasers together, and then with the three excitations sequentially. For each sample ~20 FOVs were imaged.

For the fourth color detection (Figure S6) same imaging procedure was used apart for an additional acquisition with RPA=170°.

### Single molecule FRET

For each FOV of a single smFRET species, a time-lapse of alternating 638 nm and 561 nm laser excitations operating at 100mW were taken, with 300 ms exposure per frame.

## Data analysis

### Spectral calibration

In order to enable a readout of the emission’s wavelength, a calibration between the intensity shifts in the image and the spectral information encoded within is needed. These shifts in the image domain result from the basic prisms’ dispersion curve (D_SP_(λ) in equation 1), and the RPA value as described in equation 1. In order to measure D_SP_(λ) of the prisms (since the materials of the prisms were unknown to the supplier and therefor a theoretical dispersion curve could not be calculated), images of the same 5-color bead excited sequentially with five laser excitations at 13 different RPAs, were analyzed. For each RPA, the bead’s intensity maxima per excitation source were localized, and the Y-distance from the peak of the a-chromatic image with RPA=180° was calculated. In order to reduce localization errors due to the high FRET experienced with this sample, we subtracted from each image the rescaled images of higher wavelength excitations taken with the same RPA (for example for localizing the orange peak excited by the 561 nm laser, the images of 638 nm and 785 nm excitations were rescaled to the orange emission image and subtracted from it). Each spectral window Y-distances, were then plotted and linearly fitted against sine of (180-RPA)/2 to extract the maximal dispersion values corresponding to 2D_SP_(λ) at the maximal emission wavelengths (λ_max_) (see supplemental Figure S1). Next, in order to better characterize the prism dispersion curve along the entire spectrum, we wanted to use more than 5 wavelength measurements, therefor we summed images with different excitation at the same RPA, and localized the minima between emission peaks in images of RPA≥166° were these minima were resolvable. These minima correspond with the filter’s laser line notches giving us additional 4 measurement points for the dispersion calibration curve (see supplemental Figure S2). The same linear fit to the Y-distances of minima values was performed to extract the maximal dispersion per notch wavelength (supplemental Figure S1). Next, the fitted maximal dispersion values, plotted against the wavelengths of fluorophores emission peaks and the wavelengths of filter notches, were fitted to a third order polynomial using MATLAB curve fitting tool (‘least absolute residuals’ (LAR) robust fitting method) as presented in supplemental Figure S3. The output of this fit is a calibration curve between maximal dispersion values of the CoCoS system in pixel units, and the emission wavelength. With this calibration we can transform intensity shifts in the image to spectral registration of the emission in wavelength units. Moreover, with the inverse transformation, from wavelength to pixel displacement, it is also possible to calculate the theoretical RPA values for the optimal dispersion needed in order to spectrally resolve emitters.

### Optimal RPA calculation (Table 1)

The optimal RPA values presented in Table 1 were calculated according to equation 1 using the wavelength to pixel calibration obtained in the previous section and the dyes maximal emission wavelength values obtained from their vendors websites. For dyes with emission maxima at different emission channels, the minimal spectral resolution was defined as 4 pixels separation between dispersed maxima. This separation would allow to disperse the spectra enough to achieve separation larger than the dyes PSFs (~3 pixels in diameter according to Rayleigh criterion), allowing to clearly distinguish between them and removing the need for full spectral characterization. Since color channel separation only requires small dispersion values, the PSFs themselves do not stretch considerably and only displace in position (see Table S2 in the supporting information showing that the stretching of PSF due to dispersion is usually smaller than the PSF itself), confirming that 4 pixels suffice for spectral separation. The minimal dispersion needed to spectrally resolve a pair of dyes within the same spectral window was defined as 3-pixel separation. In this case, we separate the dyes by spectral analysis which allows resolution of spectral maxima varying with only 3 pixels resolution (as was shown with CF640R and AF647 in Figure 4). For dyes with emission maxima outside of the emission channels transmission, the channel’s closest edge to the maxima was used as the emission’s maximum for calculating the minimal dispersion.

### Far-red beads spectra extraction

Spectra extraction and analysis were performed by in-house written scripts in MATLAB (MathWorks Inc.). First, in order to calculate the average spectra of the red dyes labeled beads, samples of single bead type were analyzed in the following manner. Initially, peak calling using the ‘FastPeakFind’ function^44^ was performed on the localization (180° RPA) images of multiple FOV (~8-20 depending on bead concentrations) per bead sample. Beads separated by distances shorter than the dispersion at the optimal RPA of 120° (<70×6 pixels^2^) were discarded. Next, for each bead location we searched for local maxima in the dispersed image (RPA 120°) to find for any location shifts perpendicular to the dispersion axis due to aberrations (both by the telescope lenses L3, L4 and due to the limited clear aperture of the prisms causing aberrations in large dispersion values). Next, intensity profiles were extracted from the dispersed images by taking the mean intensity values of 70×6 pixels^2^ crops around the corrected bead locations, Gaussian filtered with 1 pixel sigma (filtering was performed in order to reduce the effect of the Poisson distributed noise^16^ more evident in the dispersed image due to the lower SNR). The intensity profiles’ x-values were taken to be the displacement values from the bead location in the dispersion axis (i.e. x=0 is the location of the bead in the non-dispersed image). All profiles were then normalized between 0 and 1 and adjusted from pixel displacement values to wavelength according to the spectral calibration curve. To produce the average spectrum of a single dye used for the bead classification process, all normalized profiles originating from the same red dye beads were averaged (see supplemental Figures S9 and S10).

Optical distortion of the FOV induced by the L3 and L4 telescope lenses had created spectral shifts corresponding to beads locations in the FOV. These shifts were corrected by registering the spectral shifts compared to the median spectrum of multiple single-dye bead-samples (see supplemental Figure S14), and fitting these shifts as function of the beads spatial location to create a calibration matrix. The fitting was performed using MATLAB’s surface fit tool with local quadratic regression model (‘loess’) and least absolute residuals’ (‘LAR’) robust fit (see supplemental Figure S15). The fitted surface was later used to correct for optical aberration induced spectral shifts both in the average spectra calculation and in the dye classification procedures.

### Multi far-red dyes classification

In order to analyze spectra from FOV containing a mixture of beads labeled with different red dyes, we first performed background subtraction due to high free dyes concentration. Median pixels values taken from multi-FOV stack were used as a background mask that was subtracted from the analyzed FOV using FIJI’s^45^ ‘calculator plus’ plugin. Next, beads spectra were extracted from the background-subtracted FOV according to the procedure explained above. Spectra classification was done by cross-correlating the extracted bead spectra with the average dye specific reference spectrum obtained from FOVs containing single dye beads. The spectra were classified according to the highest cross-correlation score between the three optional spectra, with all maximal correlation scores being above 0.9, indicating good agreement. In general classification to Cy5.5 was relatively easy, with CC scores varying substantially (mean difference from the next best score was 0.35), while classification between the CF640R and AF647 is harder (mean difference from the next best score: 0.040) from obvious reasons as the spectra are less distinguishable. The spectral classification process can be also done with the theoretical spectra with similar classification results (mean difference from the next best score for Cy5.5: 0.46, CF640R and AF647: 0.047), however, with lower overall scores, probably due to pixel to wavelength calibration inaccuracies (see supplemental Figure S12).

### smFRET

The analysis of the smFRET time lapses was carried out by the iSMS software^46^ based on MATLAB (MathWorks Inc.) and additional custom MATLAB code. The analysis was performed following the standard procedure in the software’s manual. The main difference being that the two regions of interest (ROIs) selected for the dual-channel analysis were overlapped on the entire FOV which results in 3-7 times larger FRET ROI compared to standard dual-color FRET TIRF experiments. The FRET correction factors used in the analysis were calculated according to the procedure described in the recent work^15^ by Hellenkamp et al. and were the following: direct excitation (*α*) = 0.058±0.068, donor leakage (*δ*) = 0.127±0.048, *γ* factor = 0.5541±0.078 and β factor = 1.77±0.025. An elaborated account on the calculation of the various correction factors in this analysis is given in the supporting information. To assure that only a single FRET pairs were analyzed per molecule, only time traces that presented a clear single bleaching step of either the acceptor or donor upon acceptor/donor excitation (AA/ DD respectively) were taken into account in the analysis. Data points from the corrected FRET efficiency (E) and stoichiometry (S) time traces of each of the three different FRET samples (High/Medium/Low FRET) were plotted separately on 2d-histograms (see supporting Figure S16), and fitted by a Gaussian mixture model (GMM) of two 2d-gaussians to disentangle the Gaussian measurement noise (random blinking/fluctuations of the fluorophore intensities) from the actual Gaussian distribution originating from the real E and S values. The more significant 2d-gaussian distribution out of the two, was used to calculate the mean FRET efficiency value 〈E〉 for each sample (shown at the bottom of Figure 5d). These values were then used to calculate the FRET-averaged distance R〈E〉 according to the equation:

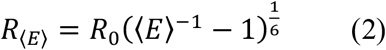

Where R_0_ is the Förster radius taken from Ref. ^15^. The FRET-averaged distance error was calculated according to the error propagation described by Ref. ^15^ taking into account the standard errors of 〈E〉 per sample, calculated from the Gaussian fits, and ΔR0 taken to be 7% of R_0_.

### NanoString

The microRNA identity corresponding to the most abundant barcode in each sample, was deduced by comparing to the results of barcode frequencies from the nCounter data (see supplemental Figure S6). The nCounter output was exported using the nSolver^TM^ analysis software (NanoString Technologies Inc.), with highest recorded abundancies of 31% (hsa-miR-150-5p), 19.3% (hsa-miR-142-3p), and 18.5% (hsa-miR-223-3p) in their respective samples. These samples were scanned with CoCoS and the most frequent barcode per sample in 10 FOVs, was assigned to the corresponding microRNA (example FOV is presented in supplemental Figure S4, example barcodes in supplemental Figure S5).

## Acknowledgments

Y.E. acknowledges support from the European Research Council Consolidator grant (Grant No. 817811), J.J. is grateful to the Azrieli Foundation for the award of an Azrieli Fellowship.

## Author contributions

J.J. and Y.E. conceived the research. J.J. designed and built the optical setup. J.J, Y.E. and Y.M designed the experiments. J.J performed all experiments and data analysis. T.C. provided the smFRET standards sample. N.S. and I.I.E. provided the NanoString samples and nCounter data. D.T designed and supervised the manufacturing of the lasers heat-sink. J.J., T.C and Y.E. wrote the manuscript. All authors read, commented on, and approved the final manuscript.

