## Supporting information for "Multi-modal Single-molecule Imaging with Continuously Controlled Spectral-resolution (CoCoS) Microscopy"

Jonathan Jeffet[1,2,3], Yael Michaeli[1,2], Dmitry Torchinsky[1,2], Ifat Israel-Elgali[4,5], Noam Shomron[4,5], Timothy D. Craggs[6], Yuval Ebenstein[1,2,3] *

1. Raymond and Beverly Sackler Faculty of Exact Sciences, Tel Aviv University, Tel Aviv 6997801, Israel
2. Center for Nanoscience and Nanotechnology, Tel Aviv University, Tel Aviv 6997801, Israel
3. Center for Light Matter Interaction, Tel Aviv University, Tel Aviv 6997801, Israel
4. Faculty of Medicine, Tel Aviv University, Tel Aviv 6997801, Israel
5. Sagol School of Neuroscience, Tel Aviv University, Tel Aviv 6997801, Israel
6. Sheffield Institute for Nucleic Acids, Department of Chemistry, University of Sheffield, Sheffield, S3 7HF UK

1. **<u>Color detection</u>**
   a. Color detection methods comparison

| **Method** | *FOV size* | *Photon loss* | *Cross talk between channels* | *Temporal channel sync.* | *# emission light paths* | *Expenses* | *Complexity* |
|---|---|---|---|---|---|---|---|
| *Split color view* | $< \frac{100\times100\ \mu m^2}{\#\ channels/\#\ cameras}$ | Passage through multiple dichroics | Dichroic and em. filter dependent | ✓ | # color channels (2-4) | $ | Complexity scales as # channels |
| *Split spectral view* | ~100-400µm² (5×30 µm²,[1] 19.2 × 19.2 µm²,[2] 120 µm²,[3] 10×10 µm²,[4]) | Passage through beam splitter | No cross-talk | ✓ | 2 | $ | Alignment: dispersive element diverts optical axis |
| *Sequential emission filter switching* | Up to ~130×130µm² | Minimal loss | Dichroic and em. filter dependent | ✗ | 1 | $ | Simple |
| *Multi cameras* | Up to ~130×130µm² per channel | Passage through multiple dichroics | Dichroic and em. filter dependent | ✓ | # color channels | $$$ | Complexity scales as # emission light paths |
| *Multi objectives and cameras* | Up to ~130×130µm² per channel | Minimal loss | No cross-talk | ✓ | # color channels | $$$$$ | Very complex alignment of multiple objectives |

*Table S1 - Summary of existing color detection methods*

b. <u>Spectral calibration</u>

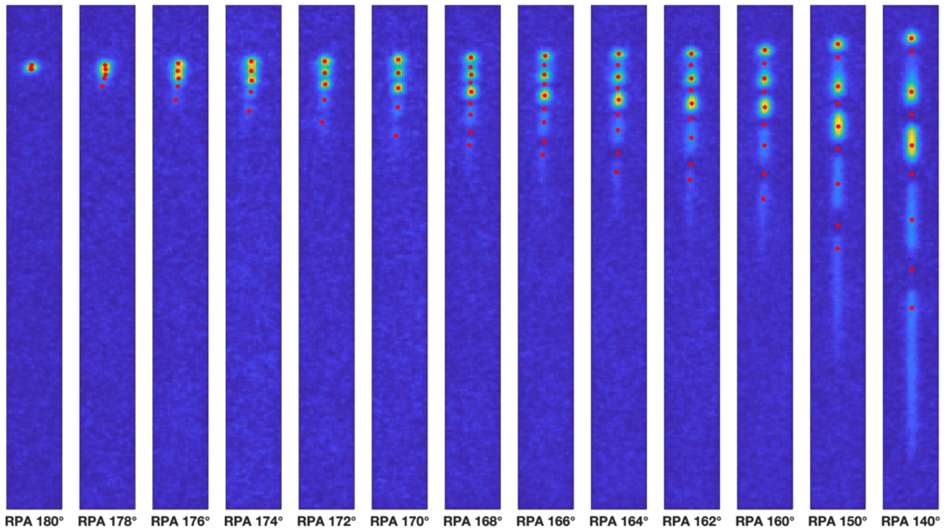

*Figure S1- Fitted extrema of an example 5-color bead spectral images at different RPAs. The red dots show the fitted extrema following the procedure explained in the methods section of the main text. The fitted Y-values of these extrema were used in the linear fit shown in Figure S2 while their fitted widths were used as error estimation.*

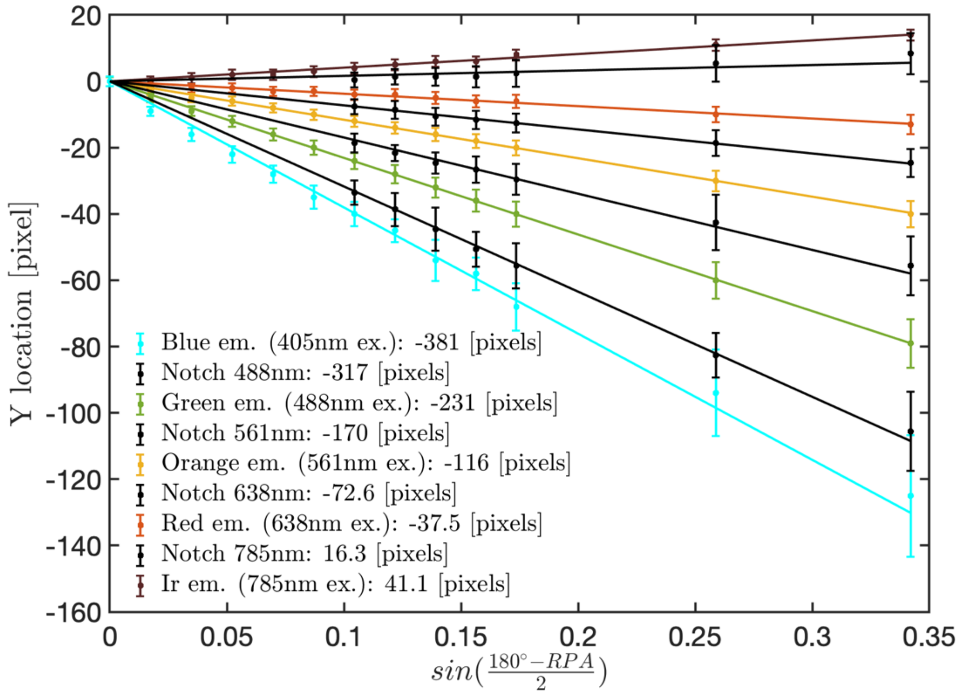

*Figure S2 – CoCoS spectral calibration using five-color silica beads. Single beads were imaged with consecutive excitation by 5 excitation sources and with varying RPAs. For each RPA the dispersed peak for each color was localized and it's Y value was plotted against the RPA according to equation (1). To gather additional information on the prism dispersion characteristics, the minima between emission peaks were also localized and assigned to the central wavelengths of the multi-notch filter. The data points were then fitted to a linear slope to deduce the maximal dispersion according to spectral window (values shown in legend). Error bars stand for the calculated width of the peaks in Figure S1.*

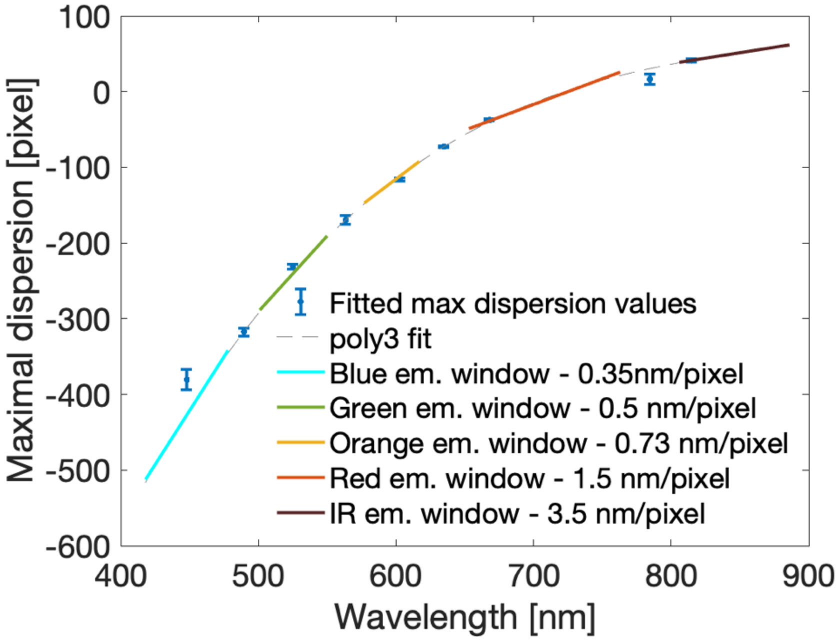

*Figure S3 - CoCoS spectral calibration using five-color silica beads. The values of maximal dispersion and their corresponding 95% confidence intervals obtained from the fit presented in Figure S2, were used to calibrate the spectral resolution of the system. A 3$^{rd}$ degree polynomial function (dashed line) was fitted to these dispersion values plotted against the emission peaks of the corresponding fluorophores. The polynomial fit was approximated to a linear function in the region of each spectral window (color coded solid lines) allowing to estimate the maximal spectral resolution of the CoCoS system in each spectral window (shown in legend) as the linear slope's reciprocal.*

c. <u>NanoString barcodes' color detection</u>

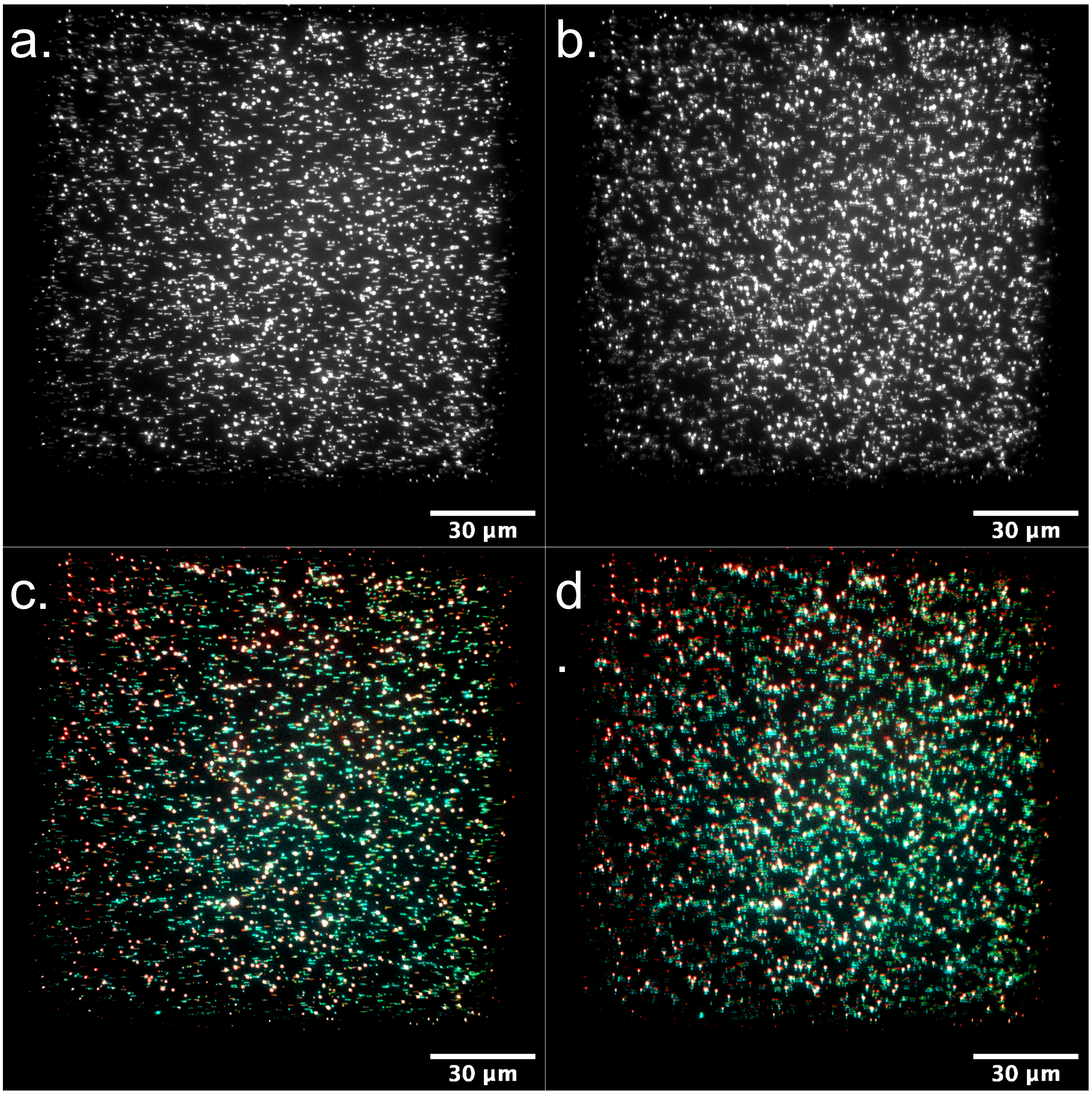

*Figure S4- Nanostring experiment full FOV with different RPA. A) Simultaneous excitation at 180° RPA with red (638mn), green (561nm) and blue (488nm) (RGB) laser sources. B) Simultaneous RGB excitation at 176° RPA. C) False colored overlay of sequential RGB excitation at 180° RPA. D) False colored overlay of sequential RGB excitation at 176° RPA. In all panels the bright spots are TetraSpeck beads used in the commercial NanoString system as fiducial markers for focus and chromatic channel registration.*

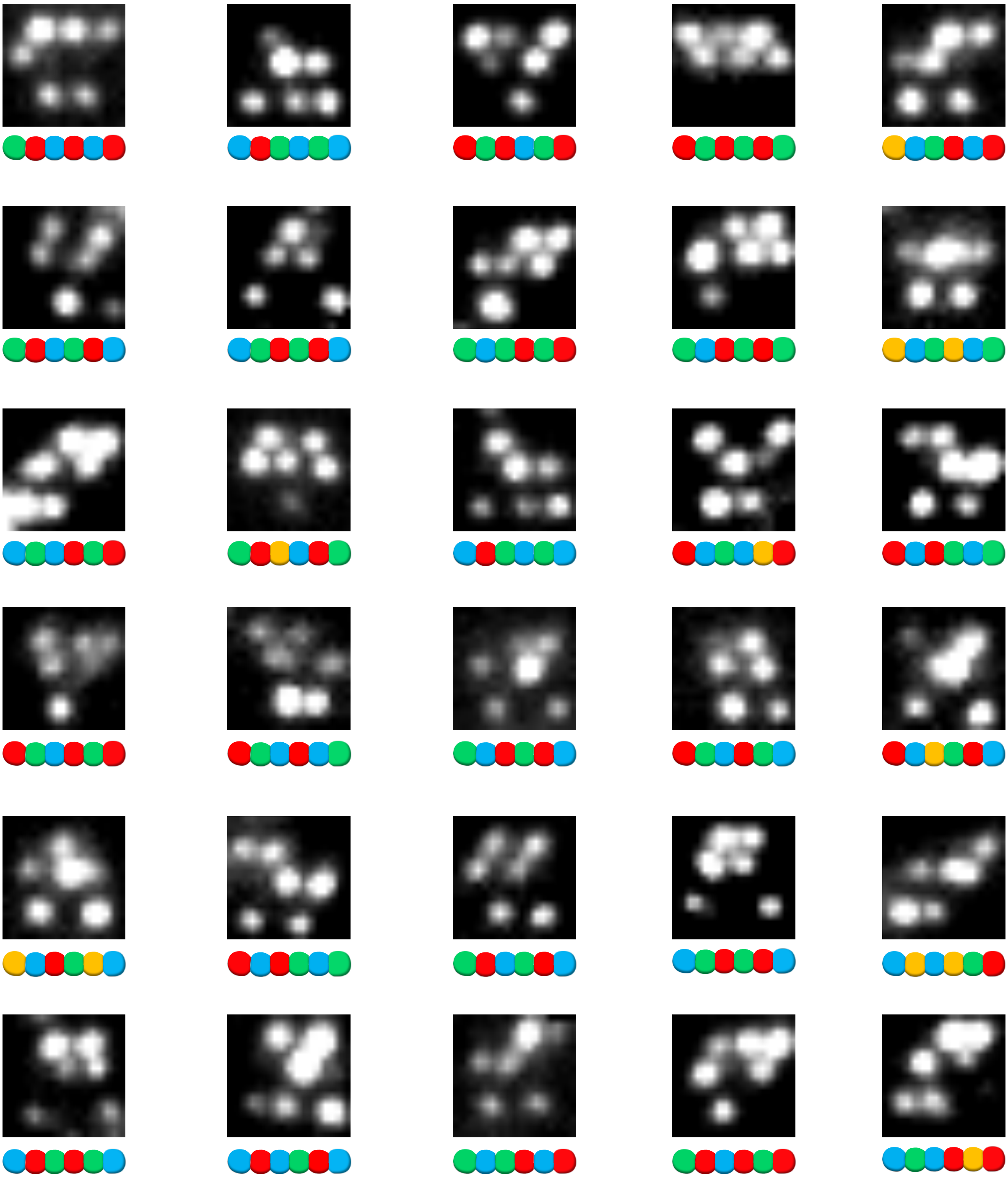

*Figure S5 – Example NanoString barcodes. 30 cropped images of different NanoString barcodes as seen in CoCoS' color detection mode with RPA=176° and simultaneous 488 nm, 561 nm and 638 nm lasers excitation. The barcodes were extracted from the FOVs of two samples. According to the nCounter barcode frequency distributions, the 26 most prevalent barcodes are shared between the three samples imaged with CoCoS. This implies that the barcodes presented here correspond to the majority of the microRNA molecules in all three samples (see Figure S6). The color representation of each barcode's dyes is presented at the bottom of each image ( blue – AF488, green – Cy3, orange – AF594, red - AF647). Differentiation between Cy3 and AF594, whose emission peaks both show in our green emission channel, was performed only when two different signals in the green channel were detected, otherwise the color code was set as green (in order to resolve the two dyes independently, smaller RPA values are needed as presented in Figure S7).*

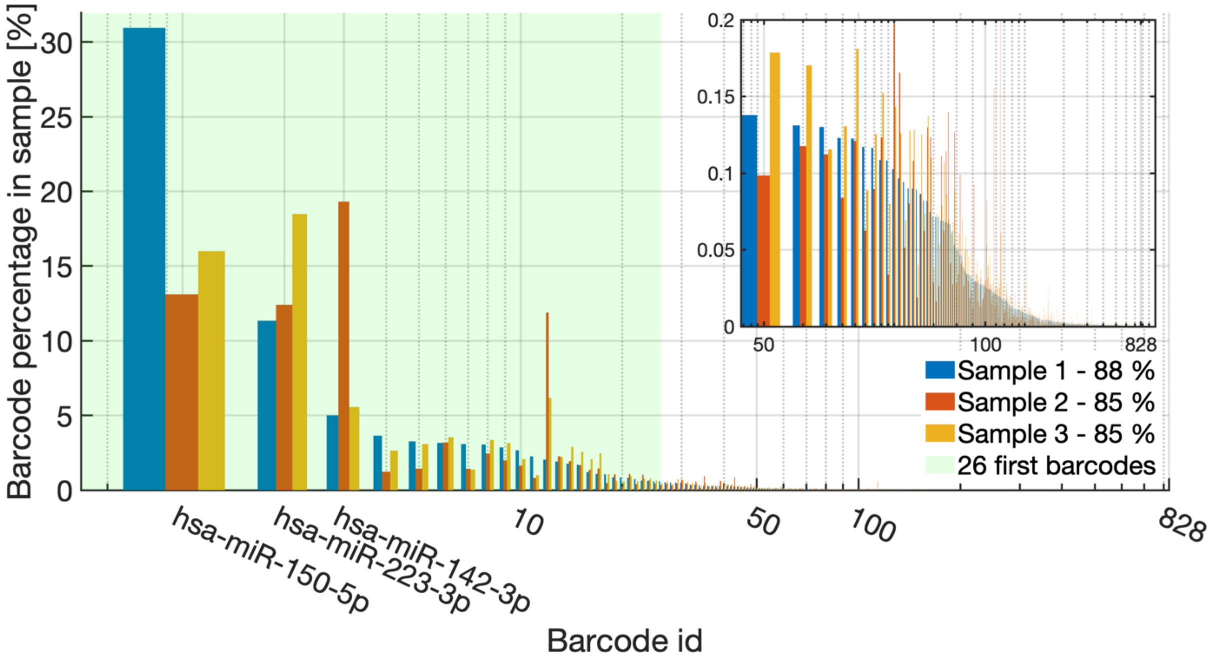

*Figure S6 - Barcode distributions in the three NanoString samples presented in the main text, obtained from the nCounter data. The most abundant barcodes in each sample are presented by their microRNA molecule name and their barcodes are shown in Figure 3. Out of overall 828 different barcodes used in these samples, the 26 most abundant barcodes (shown in green) correspond to 88% (sample 1) and 85% (sample 2 and 3) of the total microRNA molecules detected. Therefore, the barcodes displayed in Figure S5 consist the majority of the barcodes in these samples. X-axis is presented in log-scale for clearer presentation of the data, inset shows a zoom-in on the distribution of barcodes 50 to 828.*

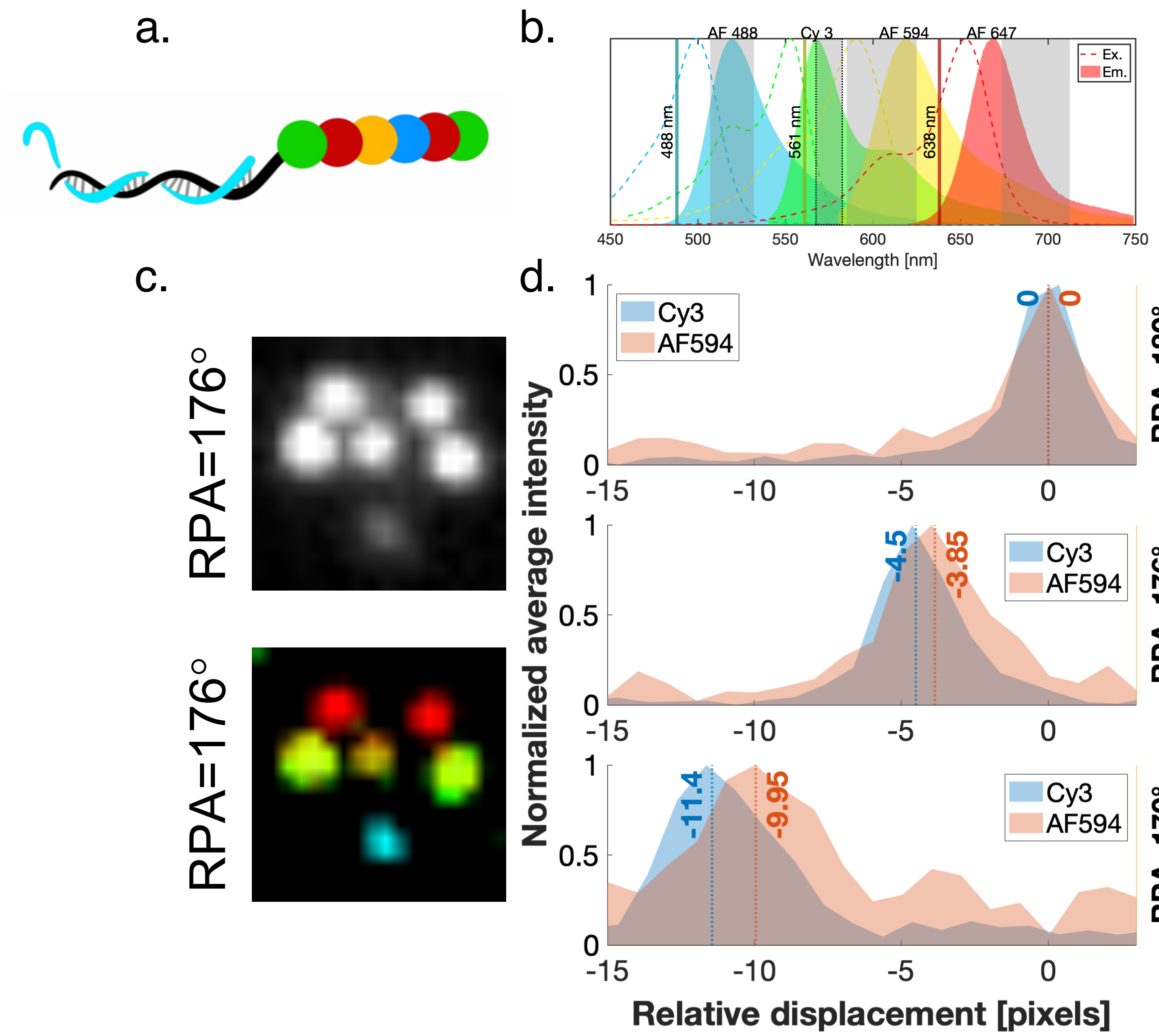

*Figure S7 – Resolving Nanostring's fourth color.* ***a)*** *illustration of a Nanostring barcode detected on the CoCoS system, and labeled with four different dyes (colors correspond to the spectra presented in* ***b****.* ***b)*** *Excitation (dotted lines) and emission (solid color) spectra of the four barcode dyes. Gray patches indicate the multi-band emission filter (MBF) channels used in this experiment, showing that the red tail of Cy3 falls within the same green channel as AF594. Dotted gray patch indicate the 575/15 nm emission filter used instead of the MBF to discriminate the Cy3 emission. The excitation lasers used in this experiment are displayed as solid lines.* ***c)*** *The barcode illustrated in* ***a****, imaged with optimal dispersion (RPA=176°). Top: single acquisition with simultaneous excitation of the three lasers) with the MBF in the emission path; Bottom: four sequential acquisitions of the same molecule, three acquisitions with the MBF and consecutive excitation with 488 nm (cyan), 561 nm (orange) and 638 nm (red) lasres andan additional acquisition with the 575/15 filter replacing the MBF in the emission path, (green) showing the Cy3 emission only.* ***d)*** *Emission intensity profiles of the same barcode (acquired from a different FOV) with 561 nm excitation and the MBF at three RPA values showing the ability to resolve different dyes with emission within the same spectral channel. The intensity peak locations at the various RPAs were caclulated by 2d gaussian fits. The intensity profiles were calculated as the mean values of a 5 pixel wide line centered at each of the peaks. All profiles were plotted relative to the peak location with no dispersion (RPA=180°). The two identicle profiles were averaged to give the Cy3 profile presented. Dotted lines show the gaussian fitted peak value relative to no dispersion (RPA=180°).*

d. PSF stretching due to minimal dispersion

| | AF 405 | AF 488 | CY 3 | ATTO 550 | AF 555 | AF 568 | AF 594 | CF 640R | AF 647 | CY 5.5 | IRDYE 800CW |
|---|---|---|---|---|---|---|---|---|---|---|---|
| **AF 405** | | 0.57 | 0.43 | 0.43 | 0.42 | 0.39 | 0.37 | 0.34 | 0.33 | 0.32 | 0.28 |
| **AF 488** | 0.57 | | 1.17 | 1.17 | 1.12 | 0.84 | 0.74 | 0.56 | 0.54 | 0.49 | 0.38 |
| **CY 3** | 0.43 | 1.17 | | | | | | 0.71 | 0.68 | 0.56 | 0.38 |
| **ATTO 550** | 0.43 | 1.17 | | | | | | 0.71 | 0.68 | 0.56 | 0.38 |
| **AF 555** | 0.42 | 1.12 | | | | | | 0.75 | 0.71 | 0.58 | 0.39 |
| **AF 568** | 0.39 | 0.84 | | | | | | 1.12 | 1.03 | 0.78 | 0.47 |
| **AF 594** | 0.37 | 0.74 | | | | | | 1.53 | 1.38 | 0.96 | 0.53 |
| **CF 640** | 0.34 | 0.56 | 0.71 | 0.71 | 0.75 | 1.12 | 1.53 | | | | 0.41 |
| **AF 647** | 0.33 | 0.54 | 0.68 | 0.68 | 0.71 | 1.03 | 1.38 | | | | 0.43 |
| **CY 5.5** | 0.32 | 0.49 | 0.56 | 0.56 | 0.58 | 0.78 | 0.96 | | | | 0.59 |
| **IRDYE 800CW** | 0.28 | 0.38 | 0.38 | 0.38 | 0.39 | 0.47 | 0.53 | 0.41 | 0.43 | 0.59 | |

*Table S2- point spread function (PSF) stretching in color detection mode (minimal RPA needed for separating two dyes). Values presented below stand for the ratio between PSF size (taken to be 3 pixels), and the dispersive stretching (in pixels) induced by the prisms (Table 1 main text). Data was calculated for a 40 nm spectral window around the emission peak of the shorter wavelength dye in each dye pair. The table shows that in most cases the stretching caused by the dispersion does not affect the PSF (values below 1) or stretch it only slightly (maximum value of 1.53 for the spectrally close dyes AF594 and CF640R).*

### 2. Single molecule Spectroscopy

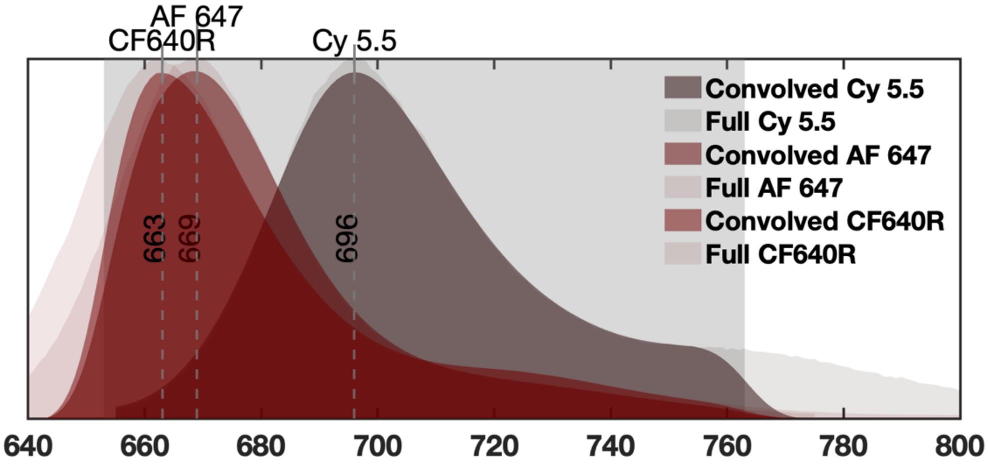

*Figure S8 – Theoretical spectra for far-red beads as seen through the CoCoS 4-notch filter. Using the available emission spectra of the different fluorophores (shown in light colors), the apparent spectra through the red emission channel (light gray rectangle) was calculated (shown in dark colors). Due to diffraction, the apparent spectra in the CoCoS system should be convolved with the PSF of the system. Therefore, a Gaussian convolution with the spectra was performed outside of the filter's transmission region. The standard deviation of the Gaussian used for the calculation was chosen according to PSF of 3 pixels in diameter and the calculated spectral resolution with RPA=120°, corresponding to the spectra presented in Figure 5 of the main text .Dotted lines indicate the convolved emission maxima with their values presented next to the lines .*

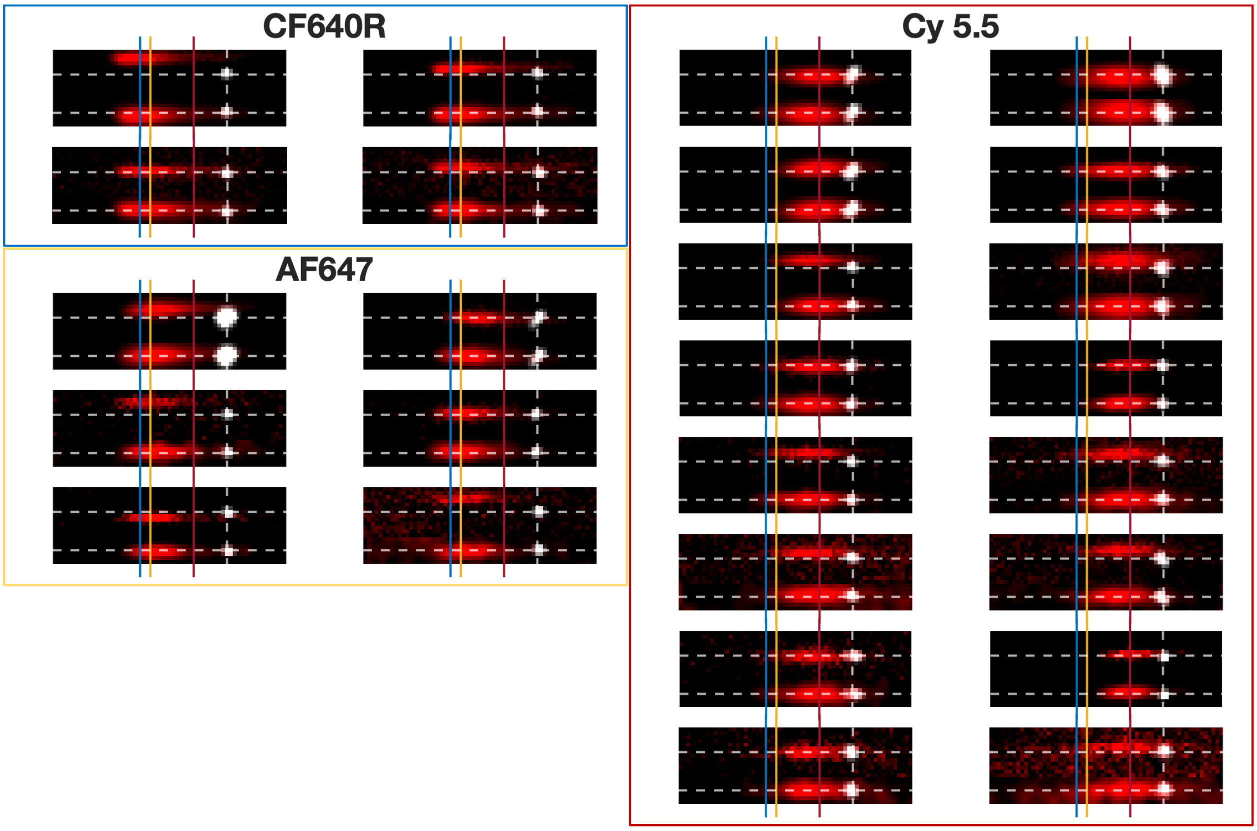

*Figure S9 – Cropped images of the single beads shown in Figure 4 in the main text compiled to their corresponding dye. Each panel shows spectral image (RPA 120°) overlaid with the localization image (RPA 180°). At the top of each panel is the raw image and at the bottom is the Gaussian filtered spectral image after perpendicular aligning and aberration correction (see Figure S15 and Figure S16). The white lines correspond to the localized coordinates of the bead, the red, yellow, and blue vertical lines show the theoretical emission maxima of Cy5.5 (696nm), AF647 (668nm) and CF640R (662nm) respectively.*

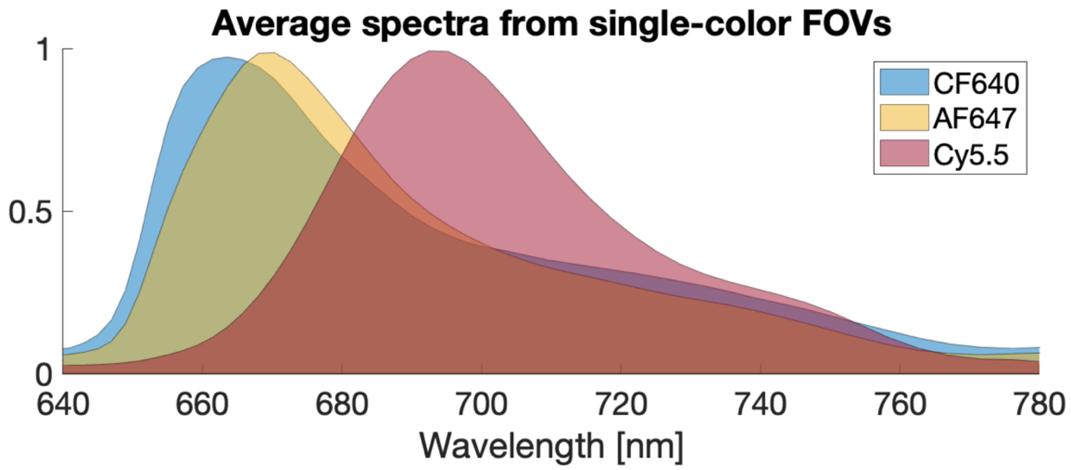

*Figure S10- Average spectra extracted from multiple FOV of single dye beads.*

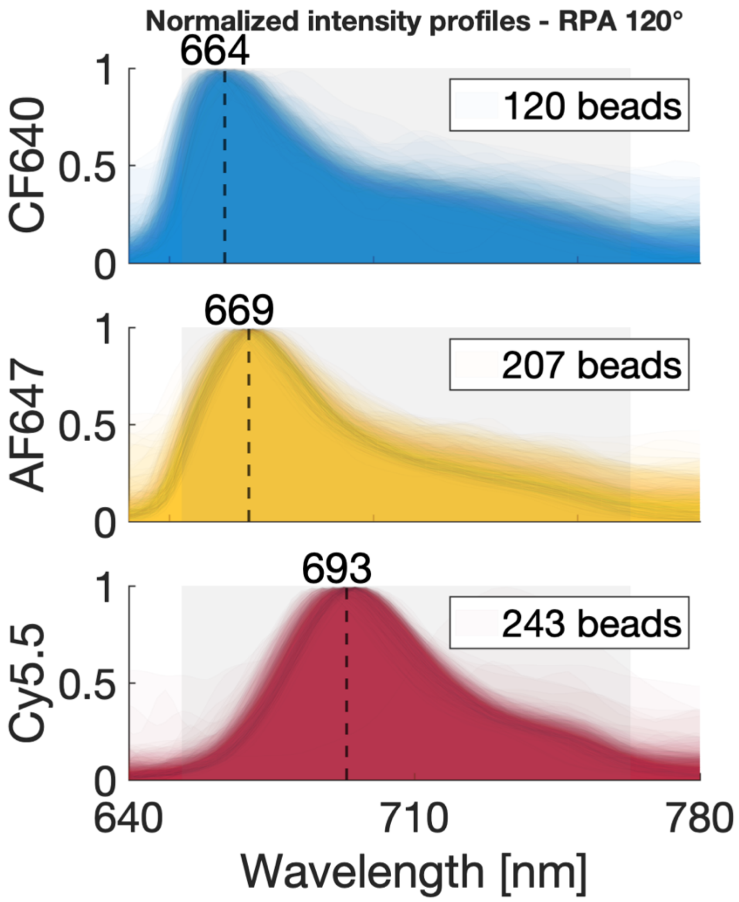

*Figure S11 - Single bead spectra used for average spectra calculation shown in Figure S10. The light gray patch corresponds with the emission filter transmission region. The number of spectra per dye is presented in the legend.*

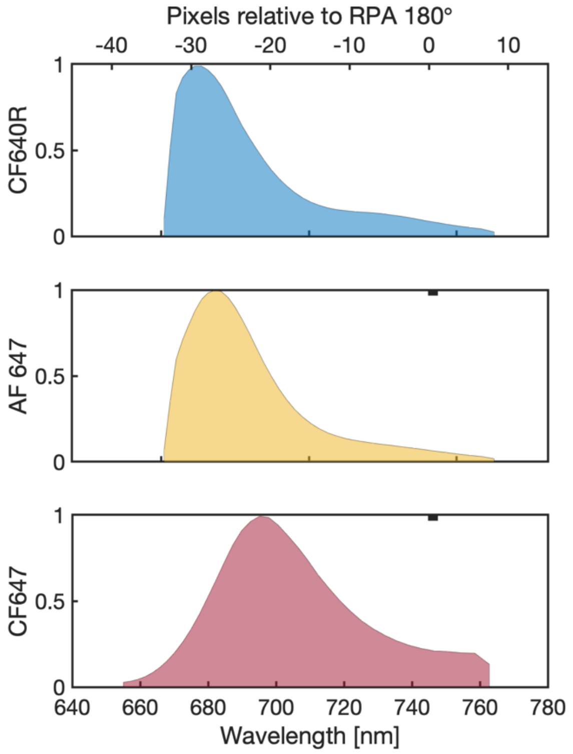

*Figure S12 - Theoretical spectra of the three dyes used in the far-red beads classification experiment. The spectra were truncated according to the red channel's emission filter, convolved with a Gaussian function to simulate the effect of point spread function spreading due to diffraction, and resampled to pixel resolution. Top X-axis shows the expected pixel distribution with RPA=120°. These spectra were used for the classification shown in Figure S13.*

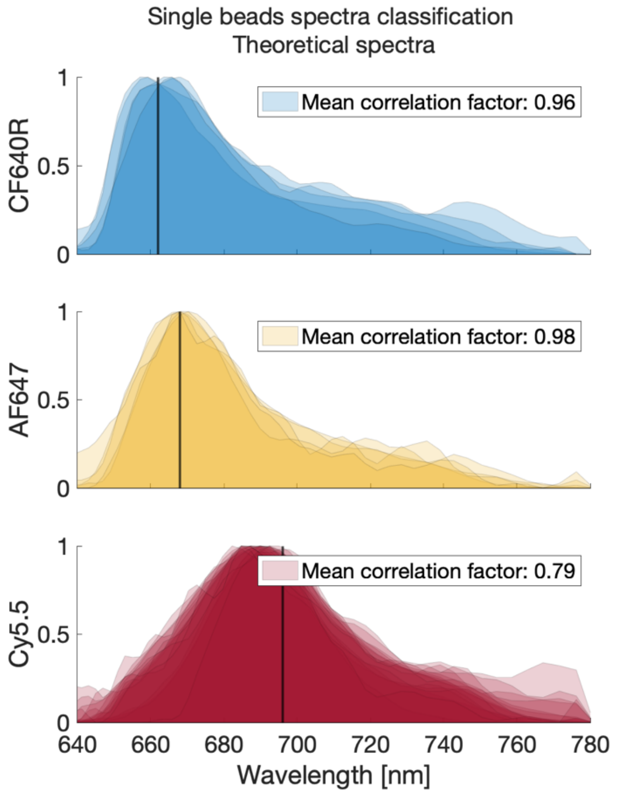

*Figure S13 - Single beads spectra from Figure 4 in the main text, classified according to theoretical spectra shown in Figure S12.*

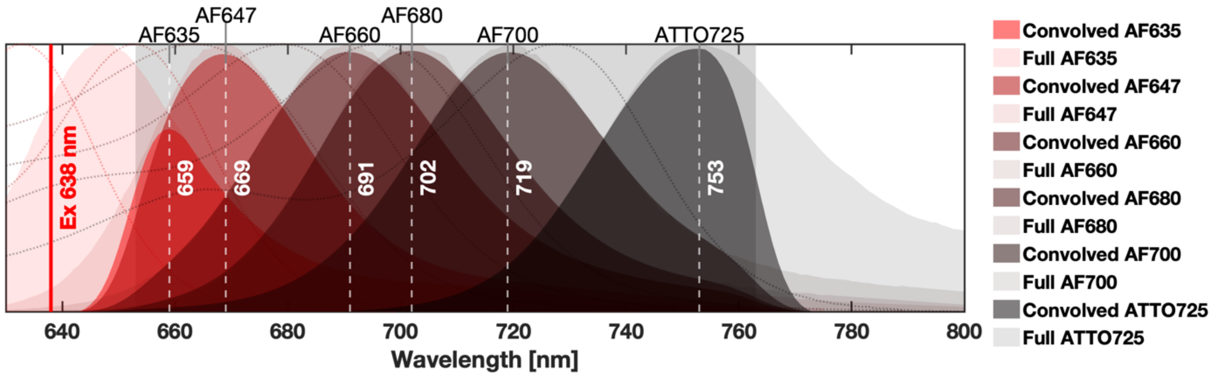

*Figure S14 – Spectra as seen in CoCoS of six commercially available far-red dyes suitable for single channel multiplexing with CoCoS. The theoretical full spectra (emission shown in solid light colors, excitation as dotted lines with corresponding colors) was convolved with gaussian by the same procedure as in Figure S8 giving the apparent spectra of the dyes (dark colors). The spectral differences between consecutive dyes is larger than the spectral difference of the CF640R and AF647 which was resolved in this work (Figure 4 main text). Dashed white lines correspond to the apparent emission peak in CoCoS.*

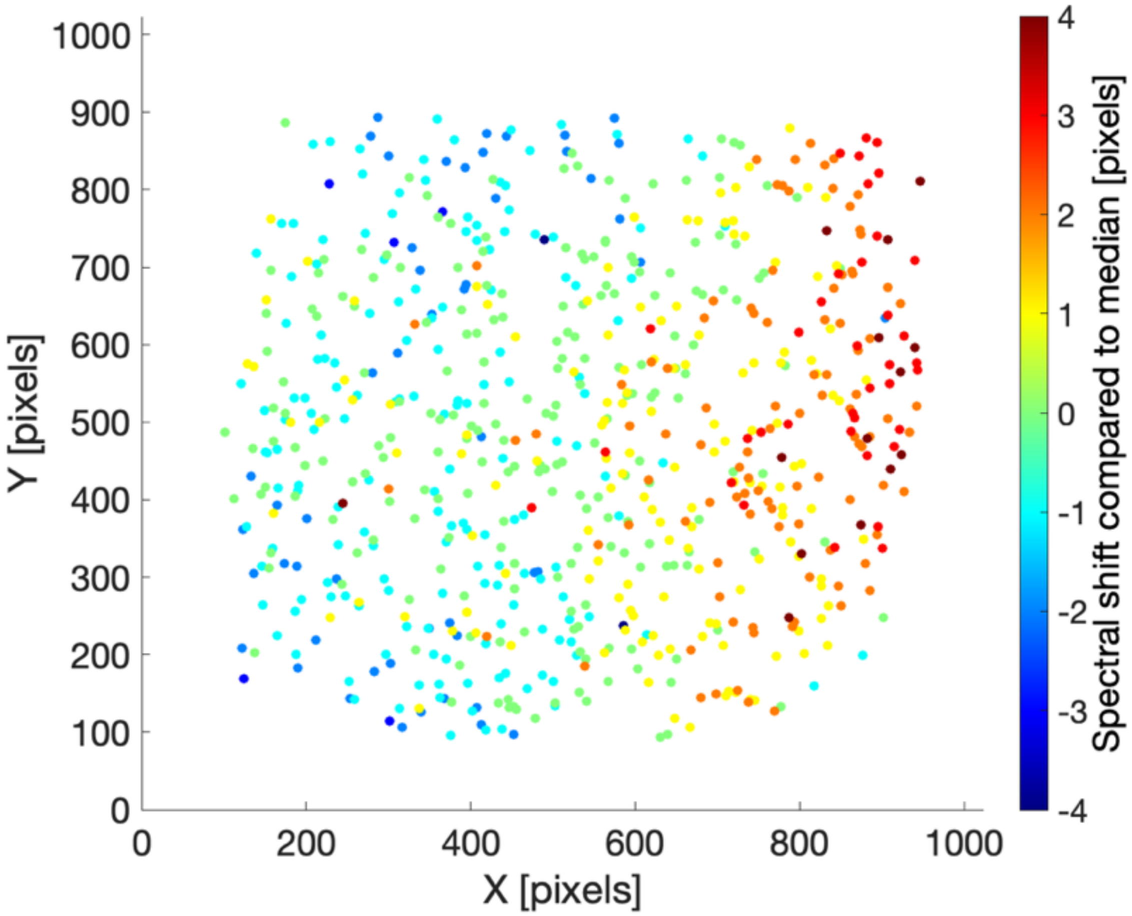

*Figure S15 – Spectral shifts as function of location in single far-red dye beads FOVs. The registered points were calculated using beads spectra from multiple single-dye samples of CF640R, AF647 and Cy5.5 beads. Color code indicates pixel shifts of emission peaks' compared to the median spectrum of all beads labeled with the same dye. The presented shifts show clear spatial dependency caused by optical aberrations induced in the imaging light path.*

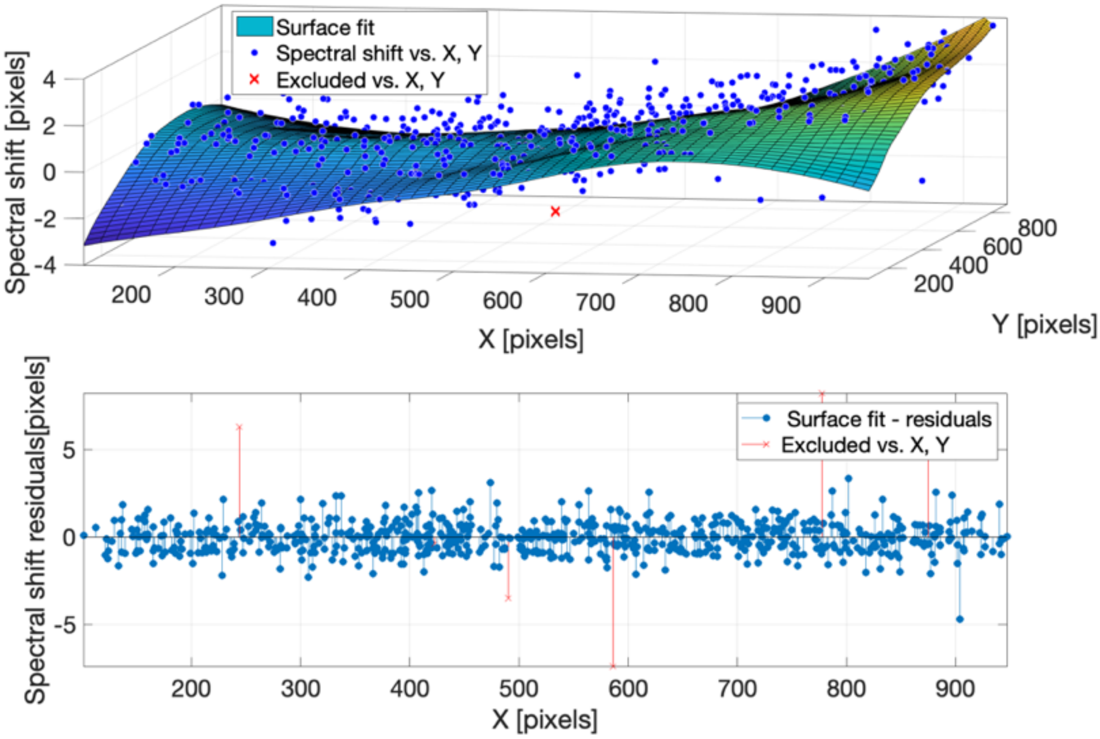

*Figure S16 – Aberration correction matrix calculation. Surface fit (top) and residual plot (bottom) to the spectral shifts presented in Figure S15. Outliers which were excluded from the fit are presented as red crosses.*

### 3. smFRET

#### a. smFRET analysis and correction factors calculation

As discussed in the main text, the analysis followed directly the standardized analysis pipeline developed by Hellenkamp et al.[5] Briefly, the procedure followed these steps: FRET traces were created with local background subtraction for all molecules in the FOV. To assure that only a single FRET pair is analyzed per molecule, only time traces that presented a clear single bleaching step in either of the acceptor/donor intensity upon acceptor/donor excitation (AA/DD respectively) traces were taken into account in the next steps of the analysis. Next, FRET efficiency (E) and stoichiometry (S) values for each time point in these traces were plotted on 2d histograms for donor only (DO) and acceptor only (AO) species from all types of samples (High, Medium and Low FRET samples). In order to calculate the global direct excitation ($\alpha$) and donor leakage ($\delta$) correction factors, a three 2d-Gaussian model was fitted to these E-S distributions (to fit the DO, AO and donor-acceptor (DA) species in each distribution). The mean ⟨E⟩ of the DO ($\langle E^{DO} \rangle$) Gaussian and the mean ⟨S⟩ of the AO Gaussian($\langle S^{AO} \rangle$ ) were used to calculate $\alpha$ and $\delta$ correction factors, according to the following equations:

$$\text{(S1)} \qquad \alpha = \frac{\langle E^{DO} \rangle}{1- \langle E^{DO} \rangle}$$

$$\text{(S2)} \qquad \delta = \frac{\langle S^{AO} \rangle}{1- \langle S^{AO} \rangle}$$

These correction factors were used to correct all E and S traces using the iSMS software.[6] Next, the $\alpha$ and $\delta$ corrected traces were used to create a new E-S histogram of DA species from the three samples taking only E-S time points prior to the first bleaching event of either donor or acceptor (see Figure S18 for full E-S histogram and Figure S19 for E-S histogram with time points prior to first bleaching event). Unaccounted acceptor or donor blinking, or partial blinking (i.e. events where blinking occurred in mid-frame resulting in intensity fluctuations), result in deviation of the data points from the main DO E-S Gaussian distribution, creating a skew towards lower FRET values. To account for this problem, each of the different samples DO E-S distributions was fitted to a 2d bi-Gaussian model distribution (see Figure S17) by a parameter-free Gaussian fit model (by the “fitgmdist” MATLAB function). Fitting the data to a bi-Gaussian model enabled to distinguish between the primary Gaussian representing the real DA FRET pairs data points, and a secondary “smeared” noise Gaussian that originated from blinking events and intensity fluctuations. The values obtained from the primary Gaussians with their standard errors were used to calculate globally the β and $\gamma$ correction factors (see Figure S20). This calculation was performed by fitting $\langle S^{DA} \rangle^{-1}$ to the corresponding $\langle E^{DA} \rangle$ values according to the following equation:

$$\text{(S3)} \qquad \langle S^{DA} \rangle^{-1} = 1 + \gamma\beta + (1-\gamma)\beta \langle E^{DA} \rangle$$

A comparison between the global β and $\gamma$ calculation vs. fitting equation (S3) to the E-S distribution is presented in Figure S21, showing that the trend of the global calculation is more appropriately following the peaks of the distribution (whereas, the direct fit misses the high FRET sample peak completely). This poor fit originates from the summation of all samples data, resulting in the accumulation of noisy data points from all three samples (due to blinking/fluorophore fluctuations), and therefor deviation of the global distribution’s center of mass towards lower FRET values. The deviated center of mass creates increased weights for the data points of lower FRET values which does not represent the actual FRET data, and therefor creates an unreliable fit as seen in Figure S21.

Finally, all FRET traces were corrected using the calculated β and $\gamma$ correction factors and the final E distributions were fitted again to a bi-Gaussian model to calculate the mean FRET efficiency per sample and its standard error (standard deviation divided by the square root of the number of data points assigned to the primary Gaussian).

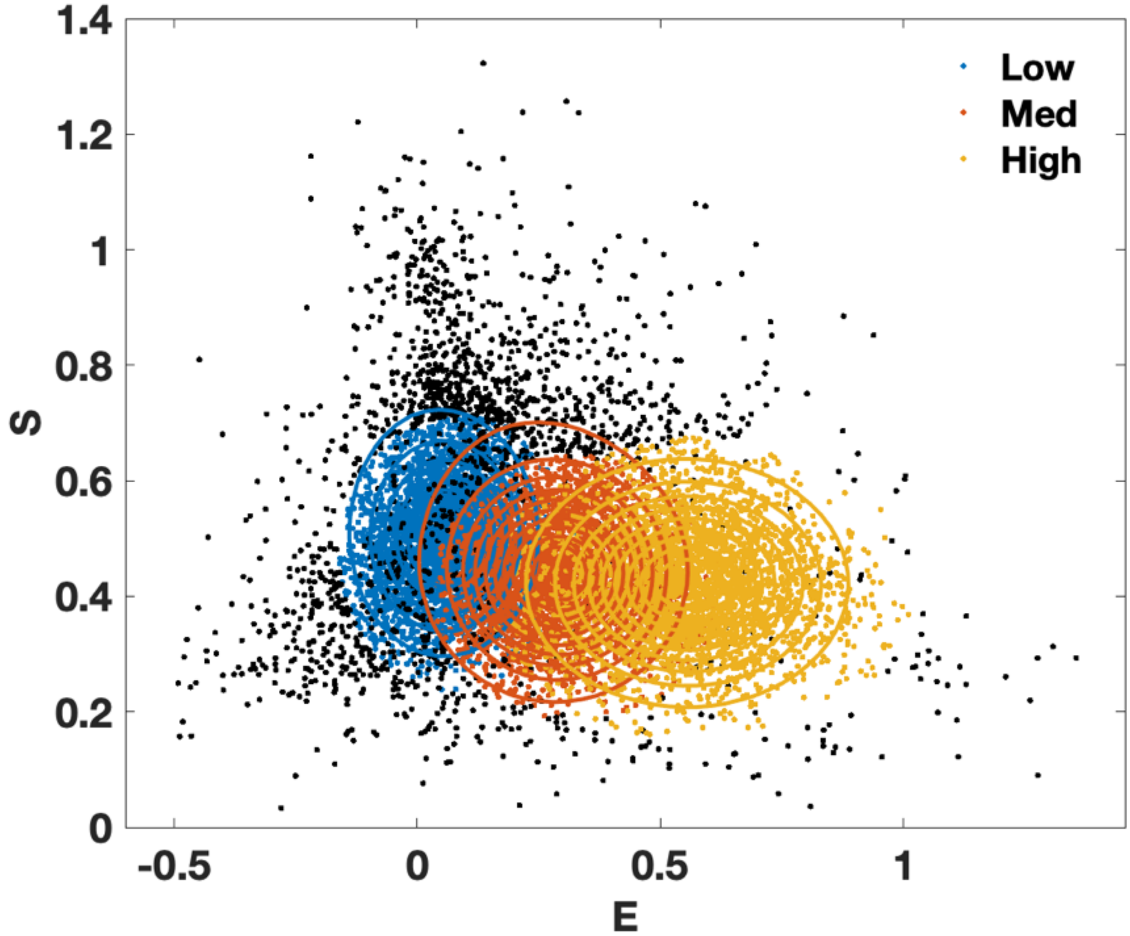

*Figure S17 – Double Gaussian fit to the different smFRET samples. For each sample the corrected FRET efficiency (E) and stoichiometry (S) histogram distribution were fitted to a parameter-free two Gaussians distribution. The two Gaussians were used to account for the measurement noise caused by blinking and intensity fluctuations. The black spots represent the data points which were classified by the model as the secondary noise Gaussian while the colored spots are the data points assigned to the primary Gaussian. The mean E and S values extracted from these fits were used for the $\gamma$ and $\beta$ correction factors calculation.*

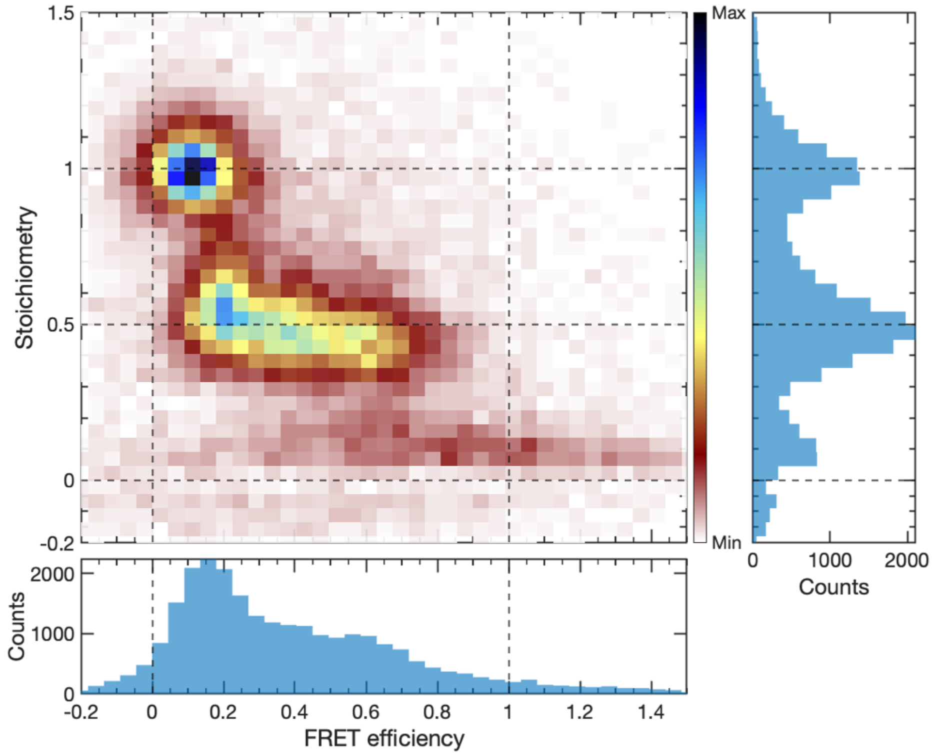

*Figure S18 – E-S histogram of all samples including donor-only and acceptor-only time points from the FRET traces, before applying any correction factors.*

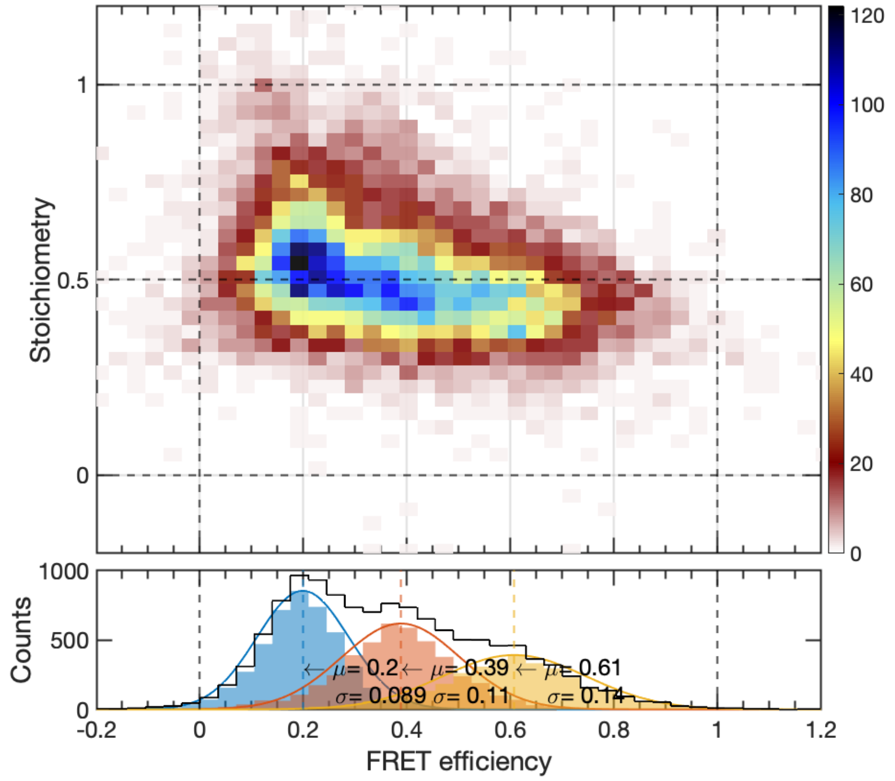

*Figure S19 - E-S histogram of all samples taking only data points prior to first bleach of either acceptor or donor and before applying any correction factors.*

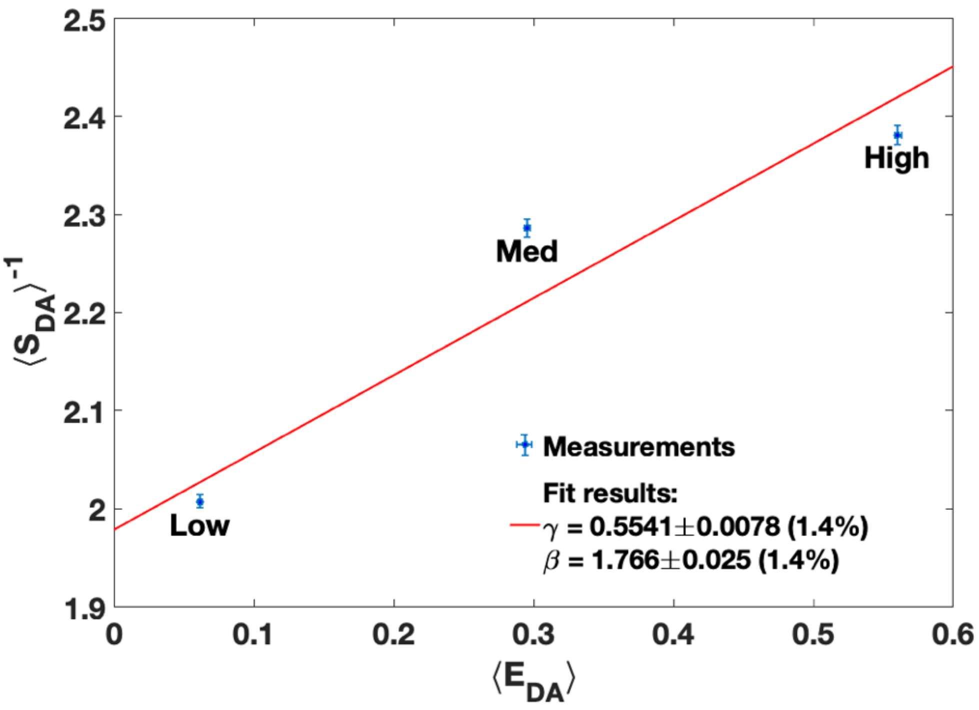

*Figure S20 – global $\gamma$ and $\beta$ correction factors calculation using linear fitting to the mean S and E values obtained from the bi-Gaussian distribution fits in Figure S17.*

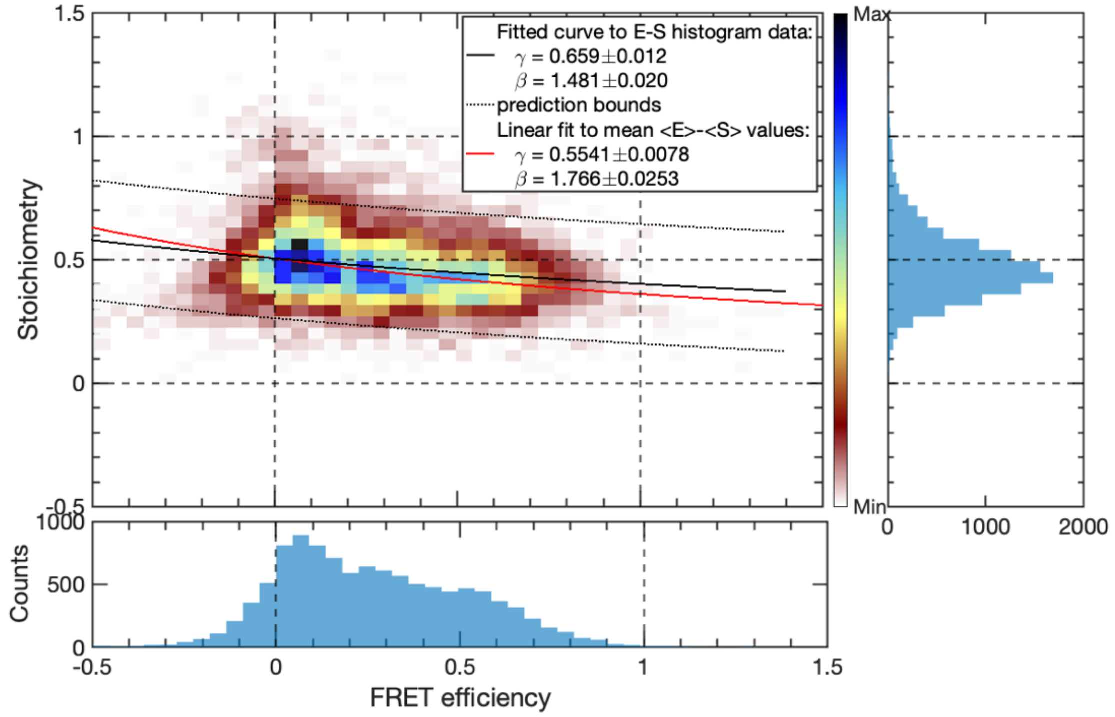

*Figure S21 - comparison between global (red) and E-S histogram curve (black) fit methods for $\gamma$ and $\beta$ correction factors calculation. The red curve follows more closely the trend of the distributions therefore this was the chosen method for correction factors calculation.*
